# Transcriptional profiling of extraocular motor neurons reveals *sim1a* as a candidate strabismus-related gene

**DOI:** 10.64898/2026.04.07.717009

**Authors:** Emily Gershowitz, Kyla Rose Hamling, Başak Rosti, Hannah Gelnaw, Grace Xiang, Cheryl Quainoo, Dena Goldblatt, Paige Leary, David Schoppik

## Abstract

Strabismus, or misalignment of the eyes, is a heritable disorder frequently associated with vision loss and decreased quality of life. Incomitant strabismus, where the degree of misalignment differs based on gaze angle, can arise from mutations in genes that regulate the development of extraocular motor neurons. To date, few such genes have been identified. The extraocular motor system is highly conserved across vertebrates, suggesting a comparative transcriptomic discovery approach would be fruitful. Using bulk and single-cell sequencing in a small accessible vertebrate, the larval zebrafish, we identified genes expressed in subpopulations of extraocular motor neurons in cranial nuclei nIII/nIV. We next assessed extraocular motor neuron number and vestibulo-ocular reflex performance after CRISPR/Cas9-mediated mutagenesis of three genes with suggestive expression patterns: *sim1a, nav2a, one-cut1*, and one known to disrupt nIII/nIV motor neuron specification: *phox2a*. Loss of *sim1a* impaired the vestibulo-ocular reflex without change to nIII/nIV motor neuron number. Our data suggest that constitutive disruptions to *sim1a* can impair nIII/nIV-dependent eye movements. More broadly, our work illuminates considerable transcriptomic diversity among extraocular motor neuron subpopulations, and establishes a pipeline to identify genes relevant to ocular motor disease etiology.

## INTRODUCTION

Strabismus, or misalignment of the eyes, is a heritable disorder of vision affecting ∼2% of the population, and is the most common childhood disorder of vision ^1^. It is frequently associated with amblyopia (vision loss resulting from inadequate visual experience during development ^2^), loss of binocular vision, and decreased quality of life in both children and adults ^3^. Incomitant strabismus, where the degree of misalignment differs based on gaze angle, can arise from mutations in genes that regulate the development of the extraocular motor system ^4,5^. While some forms of strabismus are caused by loss-of-function of known transcription factors (e.g. Duane retraction syndrome (DRS) following monoallelic *MAFB* loss of function ^6^ and congenital fibrosis of the extraocular muscles (CFEOM) resulting from biallelic mutation of *PHOX2A* ^7^), causative genes for most forms of congenital strabismus remain unknown. Recent work has highlighted several novel candidate genes relevant to ocular congenital cranial disinnervation disorders (OCCDs) ^8^, but our understanding of disease etiology is limited by a lack of insight into molecular mechanisms of extraocular motor neuron development.

The six muscles that move the vertebrate eye are controlled by six corresponding pools of motor neurons distributed across three cranial nuclei: nIII (oculomotor, 4 pools), nIV (trochlear, 1 pool), and nVI (abducens, 1 pool). Axons from each pool follow stereotyped trajectories to innervate a single muscle, and each muscle pulls the eye in a well-defined direction ^9–11^. nIII and nIV motor neurons are responsible for effecting compensatory eyes-up/eyes-down movements following vertical/torsional destabilization of the body or head. This highly conserved vestibulo-ocular reflex enables visual perception of the world to remain stable despite perturbations from self-motion. The vestibulo-ocular reflex circuit ^12,13^, including nIII and nIV motor neurons ^14^, is organized topographically by birthdate and muscle target: dorsal pools innervating the extraocular muscles that pull the eyes downward become post-mitotic earliest, while later born ventral neurons innervate the muscles responsible for upward movements. Birthdate could therefore serve as a handle to segregate functional subpopulations of extraocular motor neurons.

The larval zebrafish, a small transparent model vertebrate, offers reliable genetic access to extraocular motor neurons ^15–17^. Like all vertebrates, the zebrafish has a reliable and well-characterized vestibulo-ocular reflex ^18–22^, with comparatively simple and well-understood anatomy ^23^. Recent work in zebrafish has also demonstrated important connections between transcriptomic, anatomical, and functional characteristics of motor neuron subpopulations in the spinal cord ^24–26^. We hypothesized that a similar characterization of extraocular motor neuron transcriptional diversity might define genes that regulate development and reveal novel disease candidates.

To investigate the molecular logic that ensures proper extraocular motor neuron development, we began by performing bulk and single-cell transcriptional sequencing of extraocular motor neurons. *In-situ* hybridization experiments revealed a set of genes whose spatially- and temporally-restricted expression patterns in nIII/nIV extraocular motor neurons suggest a role in subpopulation development. Next, we evaluated nIII/nIV-dependent eye movements after constitutive loss of four candidate genes: *sim1a, phox2a, nav2a*, and *onecut1*. Loss of *sim1a* profoundly disrupted the torsional vestibulo-ocular reflex without loss of motor neurons; both nIII/nIV motor neurons and vestibuloocular reflex behavior were absent in *phox2a* mutants. Finally, *in-situ* hybridization across the vestibulo-ocular reflex circuit confirmed that *sim1a* expression was confined to nIII/nIV motor neurons. Our work reveals that *sim1a* regulates proper functional development of extraocular motor neurons. Moreover, our experiments establish a pipeline to illuminate the molecular logic that regulates extraocular motor development, a major step towards identifying and understanding the genetic determinants of incomitant strabismus.

## RESULTS

### Bulk sequencing identifies genes that label early and late-born subsets of extraocular motor neurons

Pools of motor neurons in nIII/nIV are organized topographically in both space ^27^ and across developmental time ^14^. Motor neurons that control the medial and inferior rectus muscles (MR, IR) are located in dorsal nIII, while those that control the superior rectus and inferior oblique muscles (SR, IO) are located ventrally. Dorsal nIII motor neurons are born first, followed by those in nIV (which control the superior oblique muscle, SO), and then ventral nIII Figure S1. We therefore hypothesized that comparing bulk gene expression profiles between dorsal and ventral motor neurons would yield candidate marker genes.

To isolate transcripts from early (IR/MR) and late-born (SO/SR) motor neurons, we used an optical tagging approach (Figure 1A). The transgenic line *Tg(isl1:Kaede)* ^16^ expresses the photolabile protein Kaede under control of the *isl1* promoter that labels motor neurons in three nIII pools (SR,MR,IR) and nIV (SO) ^15^. We briefly exposed larvae at 33 hours post-fertilization (hpf) to ultraviolet light, irreversibly converting the Kaede from green to red. At 33 hpf, the bulk of dorsal nIII neurons, but no ventral nIII neurons, have been born (Figure S1). We then dissected nIII/nIV from photoconverted larvae at 50 hpf, dissociated the neurons, and used fluorescence-activated cell sorting (FACS) to separate early- (red/green) and late-born (green only) neurons before RNA extraction and bulk sequencing. Almost all motor neurons are born by 50 hpf (Figure S1). We sequenced an average of 3,326 early-born and 1,577 late-born cells per replicate across four technical repeats (N=137±13 fish per repeat). Based on prior counts of ∼300 *isl1+* motor neurons per fish in nIII/nIV, we estimate a post-FACS yield of ∼10%.

**Figure 1:**
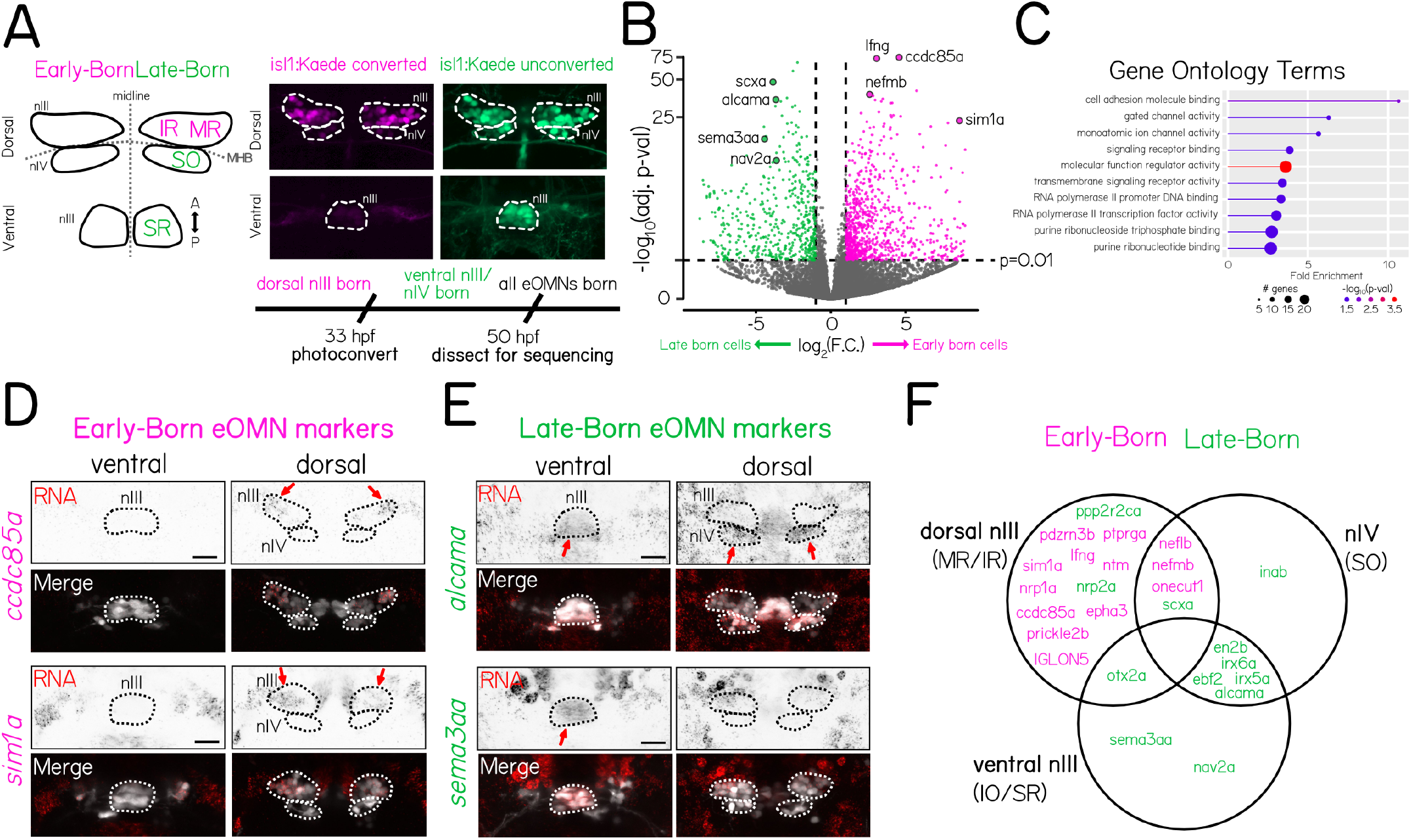
Bulk sequencing identifies genes that label early and late-born subsets of extraocular motor neurons. **(A)** (Left) Schematic of the spatial pattern of extraocular motor nuclei nIII and nIV in the larval zebrafish. Motor neuron sub-populations labeled by *isl1* (IR/MR/SO/SR) are color-coded by birthdate, with populations born before 33 hpf in magenta (“Early-Born”) and those born after 33 hpf in green (“Late-Born”). (Right) Dorsal-view confocal image of nIII/nIV motor neurons in a 50 hpf *Tg(isl1:Kaede)* larva photoconverted at 33 hpf. Only dorsal nIII neurons express photoconverted Kaede (magenta), ventral nIII and nIV neurons express only unconverted Kaede (green). **(B)** Volcano plot of gene expression fold change vs. adjusted p-value for 28,807 sequenced genes in Early-Born and Late-Born RNA libraries. Genes were significantly differentially-expressed when fold change *>* 2, and adjusted p-value *<* 0.01. 695 genes were differentially expressed, with 411 higher-expressed in early-born neurons (magenta) and 284 higher in late-born neurons (green). **(C)** PANTHER19.0 testing for over-representation of gene ontology terms for molecular function among the 695 differentially expressed genes. **(D)** Fluorescent *in-situ* hybridization examples for 2 candidate genes (*ccdc85a* and *sim1a*) predicted to be enriched in early-born OMNs. Probes targeting candidate RNA (red channel) were hybridized in *Tg(isl1:Kaede)* fish photoconverted at 33 hpf. Early-born neurons in dorsal nIII overlap with probe expression. Merge image shows the location of neurons expressing unconverted Kaede (white channel), used to draw the boundaries of nIII and nIV (dashed lines). Arrows indicate expression in dorsal nIII. Scale bar 20 µm. **(E)** Fluorescent *in-situ* hybridization examples for 2 candidate genes (*alcama* and *sema3aa*) predicted to be enriched in late-born OMNs. Probes targeting candidate RNA (red channel) were hybridized in *Tg(isl1:Kaede)* fish photoconverted at 33 hpf. Probe expression for late-born candidates is seen in ventral nIII and nIV, and does not overlap with photoconverted early-born neurons. Merge image shows the location of neurons expressing unconverted Kaede (white channel), used to draw the boundaries of nIII and nIV (dashed lines). Arrows indicate expression in ventral nIII or nIV. Scale bar 20 µm. **(F)** Venn diagram summarizing the spatial expression pattern of 25 candidate gene probes for fluorescent *in-situ* hybridization. Diagram circles indicate the subpopulations of OMNs where strongest probe expression was observed. Gene name color (magenta or green) indicates the population in which candidates were predicted to be enriched based on bulk RNA sequencing data.

Analysis of differentially-expressed genes between early and late-born motor neurons identified 695 candidate genes (411 higher-expressed in early-born neurons, 284 higher in late-born neurons) (Figure 1B). The 695 candidates included genes in functional categories implicated in neuronal differentiation and development such as cell adhesion and transcription factors (Figure 1C). Next, we used fluorescent *in situ* hybridization to evaluate the expression patterns of 25 differentially expressed genes at 50 hpf. Of the 13 genes predicted to be highly-expressed in early-born neurons, 10 were expressed exclusively in early-born neurons in dorsal nIII and the rest were expressed in both dorsal nIII and late-born nIV neurons (Figures 1D and S2). Across the candidates we evaluated, we did not see expression of genes enriched in early-born neurons in ventral nIII at 50 hpf. Of 12 genes predicted to be highly expressed in late-born neurons, 8 were expressed exclusively in late-born neurons in ventral nIII and/or nIV, 2 were expressed in both late-born and early-born neurons, and only 2 were expressed in early-born neurons (Figures 1E, 1F and S2).

We propose that differential evaluation of gene expression in early and late-born extraocular motor neurons in nIII/nIV can identify selectively-expressed transcripts that reflect temporal differences among extraocular motor neurons in nIII/nIV.

### Single-cell sequencing identifies genes expressed in sub-populations of nIII/nIV motor neurons

Temporally-defined categories broadly segregate developing neurons in nIII/nIV, but each category includes distinct pools of motor neurons. For example, late-born neurons includes motor neurons that comprise both SR and SO pools. To further validate our bulk sequencing dataset, and to refine our transcriptional classification, we performed single-cell RNA sequencing of neurons in nIII/nIV. We extracted neurons from nIII/nIV in the *Tg(isl1:GFP)* line (Figure 2A) in 48 hpf larvae. We then used FACS to isolate and place GFP^+^ neurons for plate-based CEL-Seq2^28^ transcriptomic analysis.

**Figure 2:**
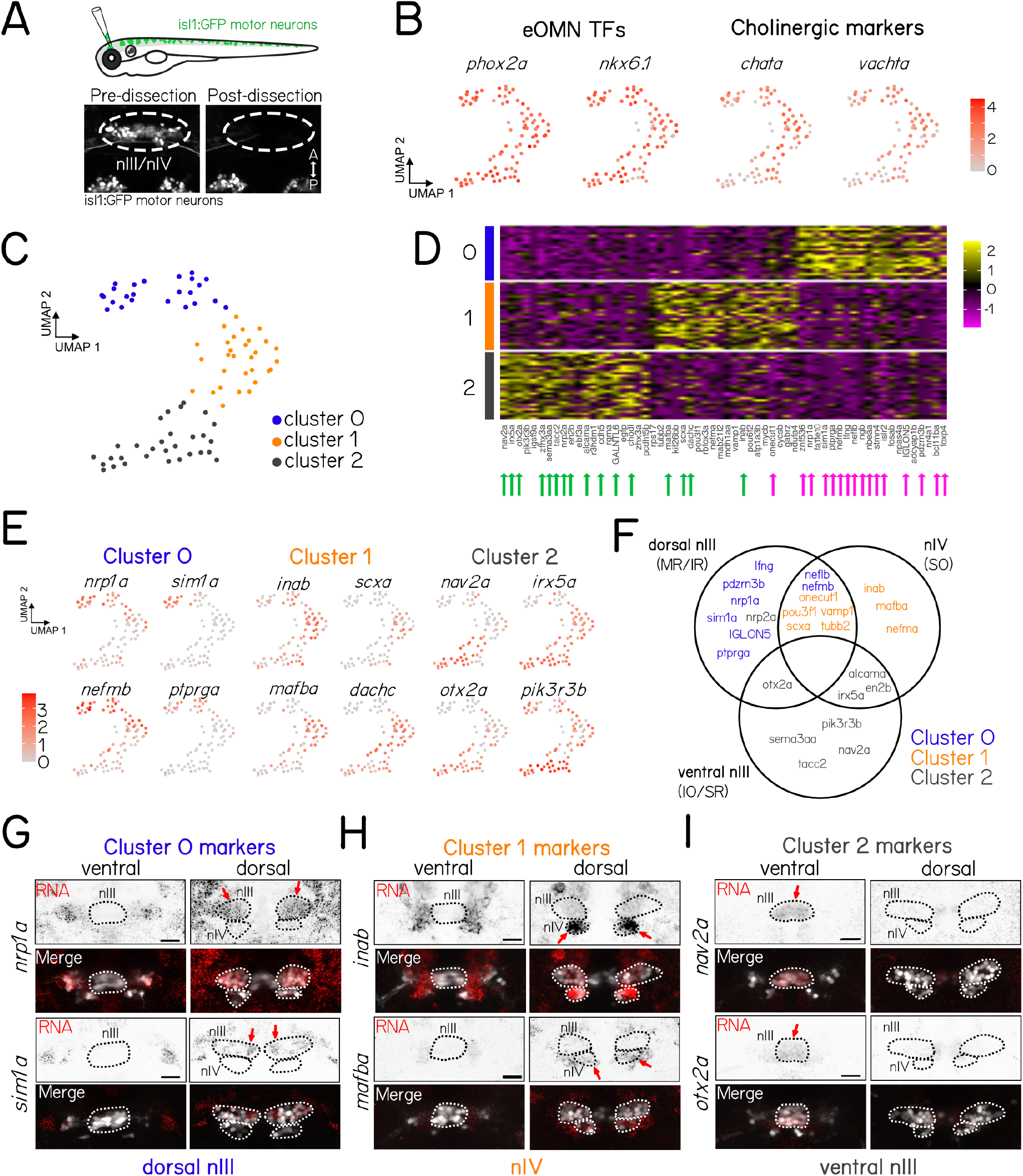
Single cell sequencing identifies genes expressed in sub-populations of nIII/nIV motor neurons. **(A)** Dissection schematic of *Tg(isl1:GFP)* ocular motor neurons (top) using targeted suction by micropipette. Example image of the same *Tg(isl1:GFP)* fish before and after targeted dissection of motor neurons in nIII and nIV show that cells of interest (dashed circle) are removed completely, while other nearby GFP+ cell populations (nV) remain intact. **(B)** Uniform Manifold Approximation and Projection (UMAP) representation of gene expression in 84 isl1:GFP+ cells dissected and sequenced at 48 hpf. Expression of canonical extraocular motor neuron gene markers show that analyzed cells have high expression of expected transcription factors (TFs) and cholinergic neuronal markers, consistent with a population of extraocular motor neurons. **(C)** UMAP representation of 84 isl1:GFP+ cells split into 3 cell clusters by gene expression profiles, representing genetic sub-populations of sequenced extraocular motor neurons. **(D)** Heat map of gene expression across all 82 cells split by UMAP cluster identify (rows) for the top 20 marker genes for each cluster (columns). Arrows below gene names indicate the marker genes that were highly expressed in late-born (green) or early-born (magenta) ocular motor neurons in previous bulk sequencing experiments. **(E)** UMAP representation of gene expression in 84 isl1:GFP+ cells for select gene markers of cluster 0, 1, and 2. **(F)** Venn diagram summarizing the spatial expression pattern of 25 cluster marker gene probes for fluorescent *in-situ* hybridization. Circles indicate the subpopulations of OMNs where strongest probe expression was observed. Gene name color indicates the UMAP cluster for which that gene was a top marker. **(G)** Fluorescent *in-situ* hybridization examples for candidate genes (*nav2a* and *otx2a*) marking cluster 0, **(H)** candidate genes (*inab* and *mafba*) marking cluster 1, **(I)** and candidate genes (*nrp1a* and *sim1a*) marking cluster 2 in single-cell sequencing experiments. Arrows indicate expression in ventral nIII (G), nIV (H), or dorsal nIII (I).

Of 192 sequenced cells, 84 passed stringent criteria for further analysis (Methods). The presence of *phox2* marks them as extraocular motor neurons in nIII/nIV (Figure 2B). Notably, extraocular motor neurons are born over an ∼28 hour period window from 22-50 hpf (Figure S1). Consequentially, some neurons in our dataset were newly-born (in ventral nIII), while others were over a day old (in dorsal nIII). This age-related gradient can be seen in the different expression levels of markers of functionally mature motor neurons such as *chata* and *vachta* (Figure 2B).

We used Uniform Manifold Approximation and Projection (UMAP) to visualize the continuum of motor neuron profiles; subsequent clustering suggests three groups with distinct RNA expression profiles (Figure 2C). The majority of the top 20 genes marking clusters 0 (12/20) and 2 (15/20) were genes we previously identified as differentially-expressed between early- and late-born categories, respectively (Figure 2D). Cluster 1 contained a small number (5/20) of genes previously identified from both early- and late-born categories, as well as many novel candidates. Qualitatively, expression levels of marker genes were consistent with either selective enrichment in a particular cluster (e.g. Cluster 0, *sim1a*, Figure 2E) or gradients of differential expression across all neurons (e.g. Cluster 2 *pik3r3b*, Figure 2E).

We repeated our fluorescent *in situ* validation on 26 of the top marker genes across the three clusters. RNA expression patterns largely matched predictions from sequencing data (Figure S2). Cluster 0 gene markers were expressed in motor neurons in dorsal nIII (Figures 2F and 2G). Cluster 1 gene markers were expressed in nIV, with three genes expressed only in nIV (*inab, mafba*, and *nefma*) and the rest expressed in nIV and dorsal nIII (Figures S2, 2F and 2H). Genes that were highly expressed in Cluster 2 (Figure 2E) were predominantly expressed in ventral nIII (Figures 2F and 2I).

Taken together, our single-cell dataset confirms our previous temporal categorization (Clusters 0 & 2 as early and late-born, respectively) and differentiates motor neurons in nIV (Cluster 1). The genes identified here are therefore candidates that might contribute to the development of selective components of the extraocular motor system.

### Select candidate genes remain subpopulation-specific across early oculomotor development

Selective gene expression patterns might define *bona fide* subpopulation-specific markers. However, the motor neurons we sequenced are different ages, and the selectivity might reflect differential sampling at specific times during development. To review: At 33 hpf, early-born dorsal nIII neurons are 0–10 hours old. By 50 hpf, all nIII and nIV neurons have been born. The oldest dorsal nIII cells are 30 hours old and have begun axogenesis, and the youngest population of ventral nIII neurons are 0–10 hours old. At 80 hpf, the oldest cells are nearing 60 hours old and the youngest motor neurons are 30 hours old; axogenesis is complete and synaptogenesis on to specific muscle targets has begun ^29^.

To determine if the subpopulation selectivity we observed persisted across early development we performed *in situ* hybridization experiments on select candidate genes at 33, 50, and 80 hpf. Both *sim1a* and *nrp1a* had persistent expression in dorsal nIII motor neurons (Figure 3A). Global expression levels for *nrp1a* varied across developmental age with particularly strong expression at 50 hpf; at this stage, expression was visible in both dorsal nIII and nIV. Expression of *nav2a* and *tacc2* persisted in ventral nIII (Figure 3B). Finally, in contrast to nIII markers, expression of *inab* and *mafba* were only strongly expressed in nIV at 50 hpf.

**Figure 3:**
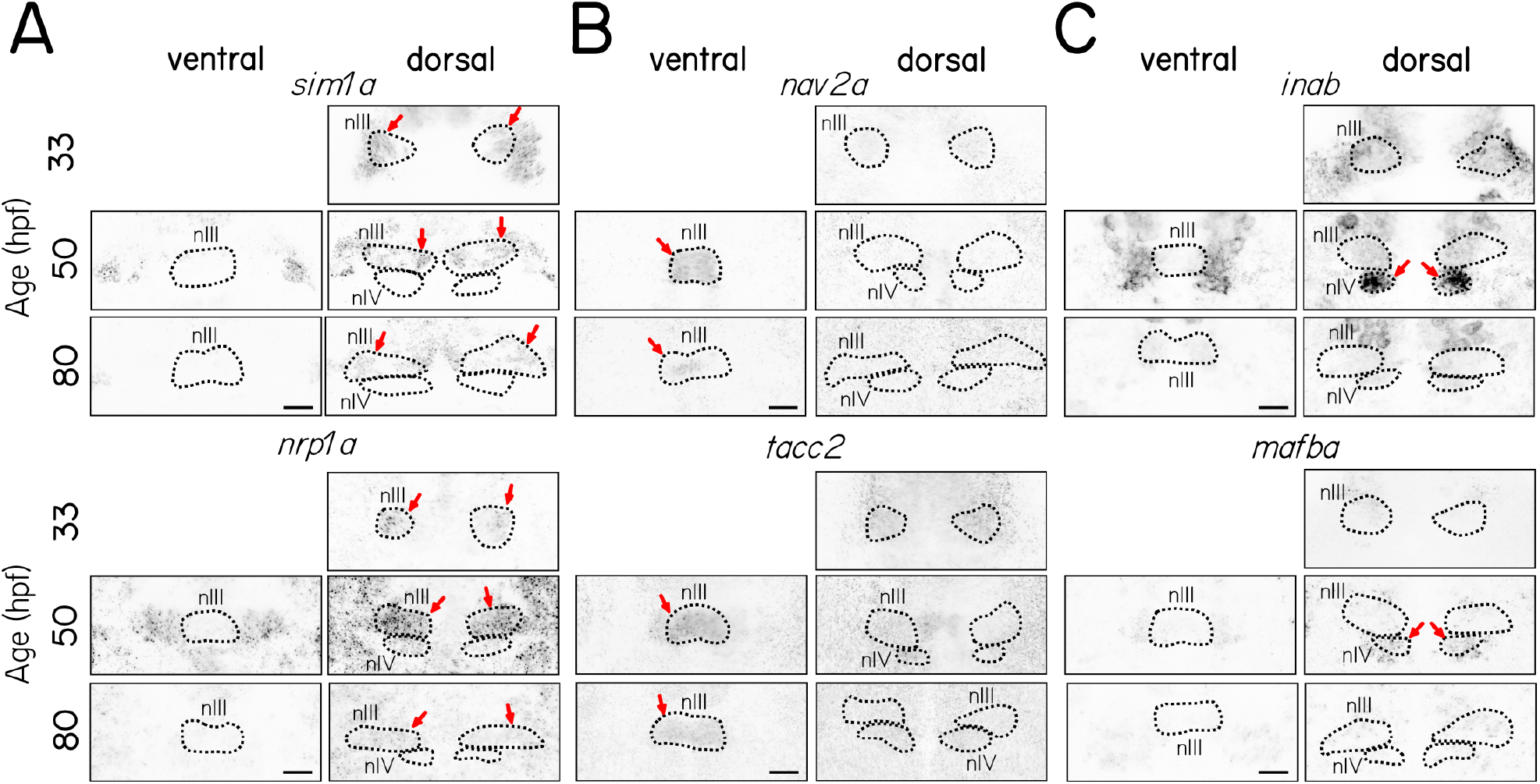
Both transiently- and persistently-expressed select candidate genes remain subtype-specific at multiple developmental stages. **(A)** Fluorescent *in situ* hybridization for *sim1a* and *nrp1a*, candidate gene markers of dorsal nIII neurons, **(B)** for *nav2a* and *tacc2*, candidate gene markers of ventral nIII neurons, **(C)** and for *inab* and *mafba*, candidate gene markers of nIV neurons, at 33, 50, and 80 hpf in ventral and dorsal nIII and nIV. Arrows indicate expression in dorsal nIII (A), ventral nIII (B) or nIV (C).

Taken together, evaluating expression at multiple timepoints suggests that our sequencing dataset contains both genes whose activity persists in specific subpopulations (e.g. *sim1a*) and genes with selective but transient expression (e.g. *inab*/*mafba*). We propose that transcriptional heterogeneity among motor neurons reflects true differences among subpopulations, rather than just differences in age.

### Subpopulation-specific gene mutations cause vertical eye rotation deficits

We next combined a genetic loss-of-function approach with an eye movement assay to evaluate what role — if any — three candidate genes play in ocular motor function. The transcription factor *sim1a* was upregulated in dorsal nIII neurons in both bulk (Figure 1D) and single-cell sequencing (Figures 2E and 2H) approaches, and remained so through early development (Figure 3A). *nav2a* was expressed in ventral nIII neurons (Figure 2F) and also remains subpopulation-specific (Figure 3B). The transcription factor *onecut1* was expressed in nIV and dorsal nIII neurons (S2B) and has been implicated in spinal motor neuron diversity ^30^. We generated loss-of-function alleles of *sim1a, nav2a*, and *onecut1* using CRISPR/Cas9 mutagenesis (Figure S3). Both *nav2a* and *onecut1* homozygous mutants were morphologically normal, but *nav2a* mutants did not survive past 7 dpf. *sim1a* homozygous mutants did not inflate their swim bladder and also did not survive past 7 dpf. Finally, we evaluated a previously validated *phox2a* loss-of-function allele that results in near-total loss of *isl1+* neurons in both nIII and nIV ^20^.

To quantify nIII/nIV-derived ocular motor function we measured the torsional vestibulo-ocular reflex following pitch (nose-up/nose-down) tilts at 6–7 dpf using a previously validated assay/apparatus ^21^. Briefly, fish were immobilized with one eye freed, placed on a rotating platform, and tilted in darkness (Figure 4A). The angle of the platform and the eye’s rotation were tracked throughout time (Figures 4B and 4C); behavioral performance is defined as the gain, or the ratio of the peak eye velocity to the peak platform velocity (35 °/sec). This vestibulo-ocular reflex can be reliably elicited after 4 dpf, but as it does not mature until 9 dpf, we expect the gain to be below 1^18,21^.

**Figure 4:**
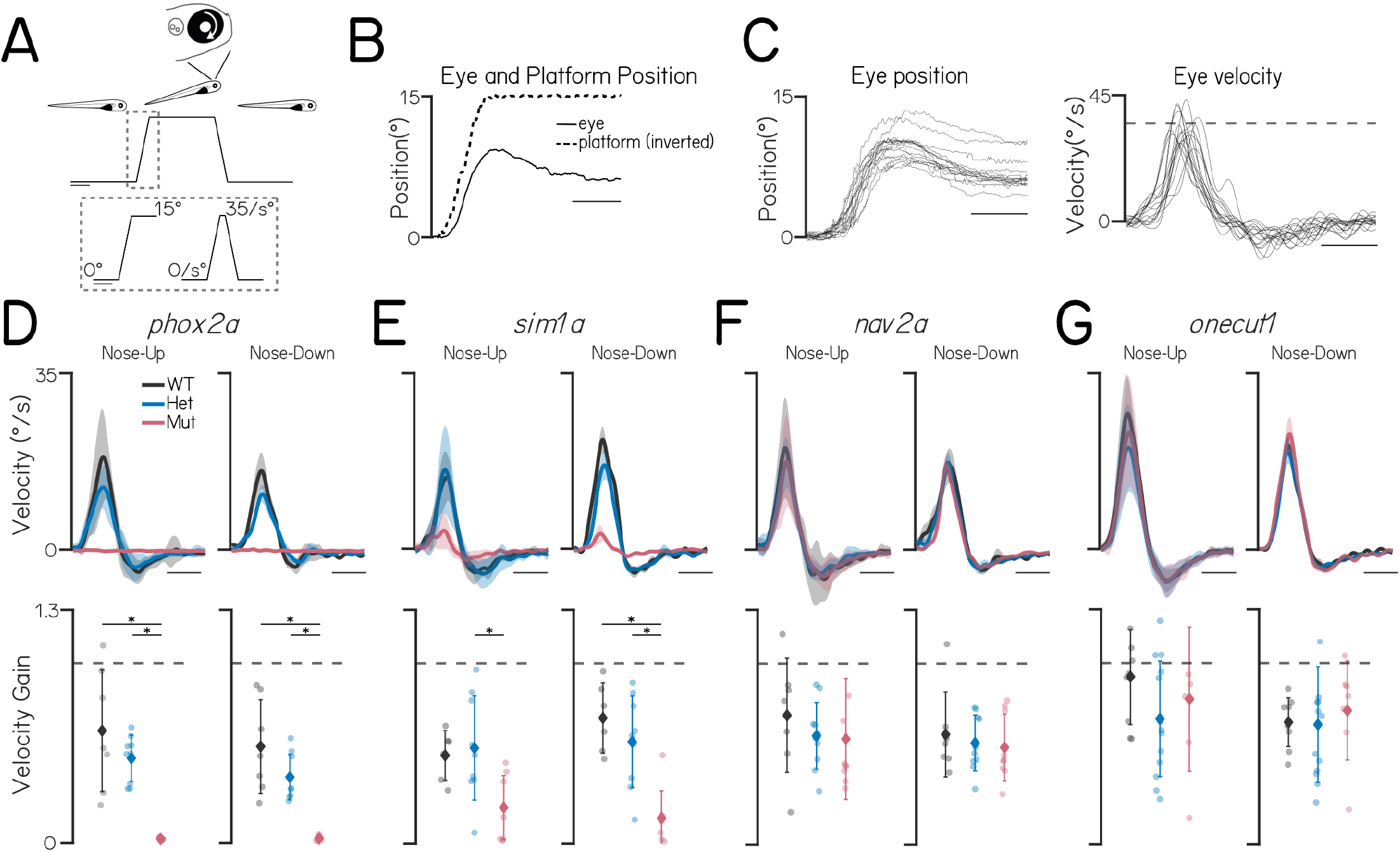
Subtype-specific gene mutations cause vertical eye rotation deficits. **(A)** Schematic of body tilt stimulus in eye rotation behavioral assay. Gray dotted box shows the trapezoidal velocity profile and angle of the tilt. Scale bars 2 seconds. **(B)** Example eye (solid) and platform (dotted) position during one stimulus step. Scale bar 0.5 seconds. **(C)** Left: eye position of a 7 dpf wildtype larva during each 15° nose-up tilt step of an experiment. Right: eye velocity of the same fish during nose-up tilts. Dashed gray line at the peak tilt velocity (35°/s). Scale bar 0.5 seconds. **(D)** Top: Average eye velocity ± SEM of wildtype siblings (WT, gray), *phox2a* heterozygotes (Het, blue), and *phox2a* homozygous mutants (Mut, pink) (n= 7, 9, and 6 fish) in response to pitch tilts in the nose-up (left) and nose-down (right) direction. Scale bar 0.5 seconds. Bottom: Gain (peak eye velocity / peak platform velocity) for each fish during nose-up (left) and nose-down (right) tilts. Wildtype versus mutant nose-up gains, P_Tukey’s_ = 1.4 *×* 10^-4^; heterozygote versus mutant nose-up gains, P_Tukey’s_ = 0.0017; wildtype versus mutant nose-down gains, P_Tukey’s_ = 8.1 × 10^-5^; heterozygote versus mutant nose-down gains, P_Tukey’s_ = 0.025. Color indicates genotype as above. Gray dashed line at gain = 1 **(E)** Top: Average eye velocity ± SEM of wildtype siblings (gray), *sim1a* heterozygotes (blue), and *sim1a* homozygous mutants (pink) (n= 6, 9, and 8 fish). Scale bar 0.5 seconds. Bottom: Gain for each fish. Wildtype versus mutant nose-up gains, P_Tukey’s_ = 0.063; heterozygote versus mutant nose-up gains, P_Tukey’s_ = 0.016; wildtype versus mutant nose-down gains, P_Tukey’s_ = 2.2 × 10^-4^; heterozygote versus mutant nose-down gains, P_Tukey’s_ = 0.0013. **(F)** Top: Average eye velocity ± SEM of wildtype siblings (gray), *nav2a* heterozygotes (blue), and *nav2a* homozygous mutants (pink) (n= 7, 10, and 9 fish). Scale bar 0.5 seconds. Bottom: Gain for each fish. Wildtype versus mutant nose-up gains, P_Tukey’s_ = 0.63; heterozygote versus mut nose-up gains, P_Tukey’s_ = 0.99; wildtype versus mutant nose-down gains, P_Tukey’s_ = 0.73; heterozygote versus mutant nose-down gains, P_Tukey’s_ = 0.96. **(G)** Top: Average eye velocity ± SEM of wildtype siblings (gray), *onecut1* heterozygotes (blue), and *onecut1* homozygous mutants (pink) (n= 9, 13, and 7 fish). Scale bar 0.5 seconds. Bottom: Gain for each fish. Wildtype versus mutant nose-up gains, P_Tukey’s_ = 0.73; heterozygote versus mutant nose-up gains, P_Tukey’s_ = 0.75; wildtype versus mutant nose-down gains, P_Tukey’s_ = 0.88; heterozygote versus mut nose-down gains, P_Tukey’s_ = 0.81.

As expected, *phox2a* homozygotes did not move their eyes in response to either nose-up or nose-down pitch tilts (Figure 4D). Similarly, the vestibulo-ocular reflex gain was markedly reduced in *sim1a* homozygotes (Figure 4E). In contrast, gain was indistinguishable between wild-type / heterozygous siblings and larvae with mutations in either *nav2a* or *onecut1* (Figures 4F and 4G). We propose that *sim1a* is indispensable for a proper nIII/nIV-dependent vestibulo-ocular reflex.

The vestibulo-ocular reflex circuit consists of sensory afferents in the stato-acoustic ganglion, central projection neurons in the tangential nucleus and motor neurons in the oculomotor and trochlear nuclei (Figure 5A). Defects in any of these areas could compromise the vestibulo-ocular reflex. *phox2a* expression was only observed in nIII/nIV (Figure 5B), and *phox2a* mutants have few or no *isl1+* oculomotor/trochlear cells (Figure 5C). In contrast, we observed no differences in the number of *isl1+* motor neurons in nIII and nIV in any of our novel mutant alleles (Figures 5D to 5F). Finally, *sim1a* was not expressed in either the stato-acoustic ganglion or the tangential nucleus at 33, 50, and 80 hpf (Figure 5G). Our data suggests that *sim1a* does not specify, but instead regulates proper functional development of nIII/nIV extraocular motor neurons.

**Figure 5:**
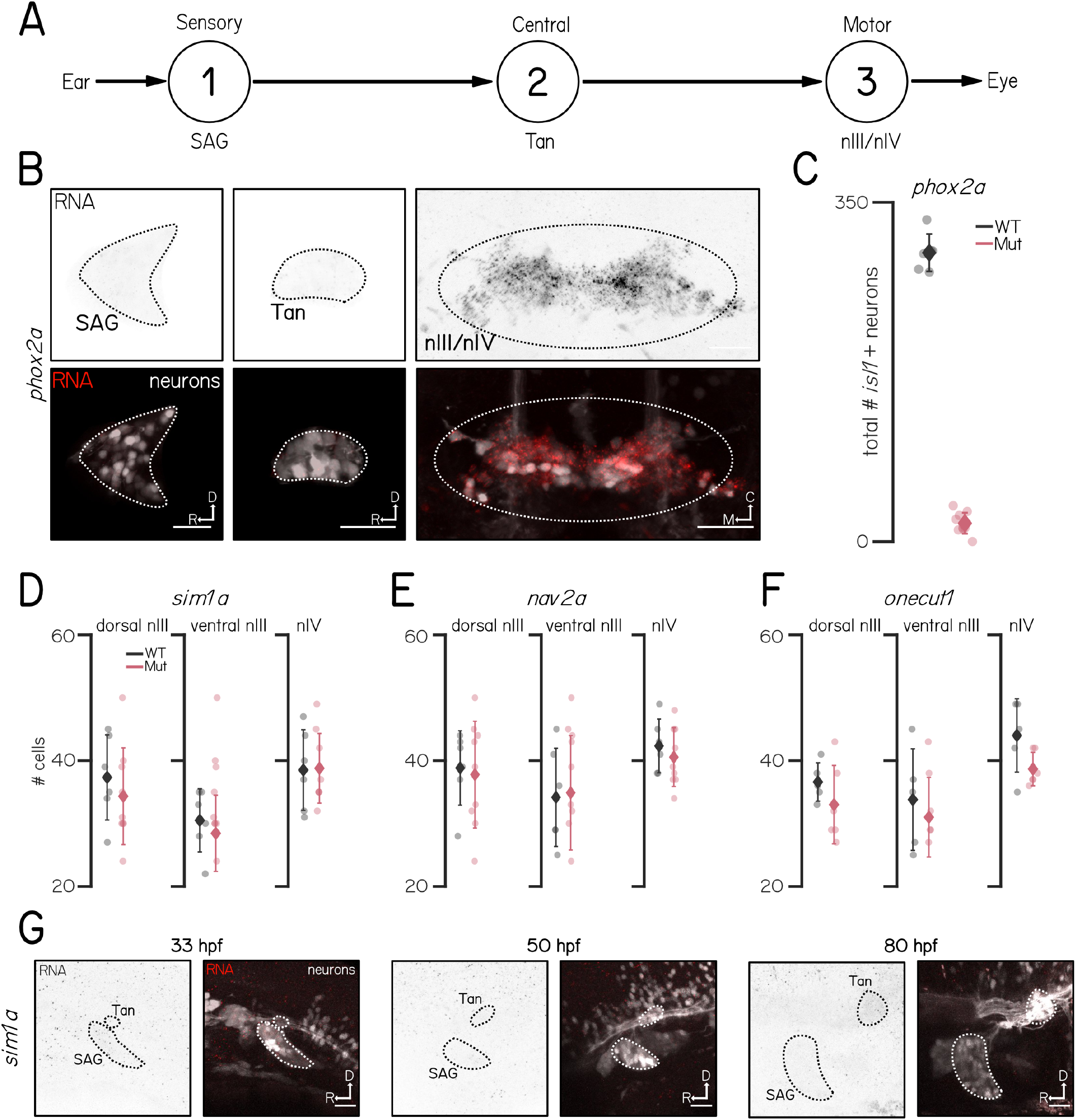
*phox2a* and *sim1a* phenotypes likely reflect impairments to motor neurons. **(A)** Schematic of vestibulo-ocular reflex circuitry. **(B)** Fluorescent *in situ* hybridization for *phox2a* in the stato-acoustic ganglion (1, left), tangential nucleus (2, center), and oculomotor and trochlear nuclei (3, right) at 5 dpf. Scale bars 25 µm. Data from ^20^. **(C)** Total number of *isl1+* cells in nIII/nIV of *phox2a* wildtype control siblings (gray) and homozygous mutants (pink). Data from ^20^. **(D)** Number of *isl1+* cells in dorsal nIII, ventral nIII, and nIV of *sim1a* wildtype control siblings (gray) and homozygous mutants (pink). **(E)** Number of *isl1+* cells in dorsal nIII, ventral nIII, and nIV of *nav2a* wildtype control siblings (gray) and homozygous mutants (pink). **(F)** Number of *isl1+* cells in dorsal nIII, ventral nIII, and nIV of *onecut1* wildtype control siblings (gray) and homozygous mutants (pink). **(G)** Lateral view of fluorescent *in situ* hybridization for *sim1a* in the stato-acoustic ganglion (SAG, 1) and tangential nucleus (Tan, 2) at 33 (left), 50 (center), and 80 hpf (right). Scale bars 25 µm.

## DISCUSSION

We measured transcriptional diversity between subpopulations of extraocular motor neurons and discovered a novel role for *sim1a* in ocular motor development. Validated bulk sequencing identified markers of early- and late-born subsets of extraocular motor neurons in the oculomotor and trochlear nuclei. Next, single-cell sequencing and fluorescent *in-situ* hybridization experiments revealed differentially-expressed genes marking specific subpopulations that remain subpopulation-specific through early development. Finally, we used the vestibulo-ocular reflex to evaluate loss-of-function phenotypes for four candidate genes. Of these, *sim1a*, a novel candidate expressed in early-born dorsal nIII neurons, is necessary for proper tilt-evoked eye movements. We propose that our pipeline from transcriptomics to behavior can discover and define roles for candidate genes responsible for extraocular motor neuron development. This work is therefore a key step towards understanding the etiology of congenital incomitant strabismus.

### Loss of *sim1a* disrupts the vestibulo-ocular reflex

*SIM1*, after *Drosophila single minded (sim)*, is a basic helix-loop-helix transcription factor ^31^ that is important for neural development in flies ^32^, mice ^33^, and humans ^34^. In zebrafish, *sim1a* has also been implicated in the development of the kidney and associated cells ^35^, acting downstream of retinoic acid ^36^ and Notch/Tbx2a ^37^ signaling pathways to specify cell fate. Similarly, *sim1a* specifies diencephalic dopaminergic neuron ^38^, hypothalamic neuron ^39–41^, and V3 spinal interneuron fates, the latter depending on exposure to Notch ^42^. In the hypothalamus, *sim1a* interactions with its binding partner *arnt2* regulate guidance of hypothalamo-spinal axons by interfering with Robo2-mediated repulsion ^43^. Prior work thus establishes *sim1a* as an excellent candidate to specify fate and/or regulate axonal projections of extraocular motor neurons in nIII.

What role might *sim1a* play in the development of nIII motor neurons? We did not observe any differences in the number of nIII isl1:GFP^+^ motor neurons in *sim1a* mutants, and we see partial preservation of both up/down eye movements. Together, these findings suggest aberrant axonal guidance or functional disruptions rather than a fate change after loss of *sim1a*. Intriguingly, in mice, loss of *Sim1* disrupts the dorsal clustering and electrophysiological diversification of early-born, but not late-born, V3 spinal interneurons ^44^. Future experiments in *sim1a* mutants characterizing nIII nerve branching/muscle innervation ^29^ and functional responses of dorsal nIII motor neurons to vestibular stimuli ^45^ will speak to the anatomical and functional contributions of *sim1a*.

*sim1a* expression is restricted to dorsal nIII, which contains inferior rectus motor neurons that move the eyes down after nose-up pitch tilts. However, *sim1a* mutants show markedly reduced vestibulo-ocular reflex gains in both directions on the pitch axis. While several cases of deletions on the long arm of chromosome 6, which contains the human *SIM1*, present with strabismus ^46–48^, (though see ^49^) comprehensive characterization of the strabismus from these patients is not available. The absence of *sim1a* expression elsewhere in the vestibulo-ocular reflex circuit argues against an upstream deficit in sensory/central processing of tilts. Notably, humans show similarly “paradoxical” vestibulo-ocular reflex impairments in cases of fourth nerve palsy in adulthood, thought to reflect adaptive plasticity ^50^. Alternatively, impaired basal tension of extraocular muscles would shift the eyes’ resting orientation, disrupting both up and down rotations. Future experiments could measure visually-evoked vertical eye movements ^22^ and/or medial rectus-dependent horizontal eye movements ^51^ in *sim1a* mutants to better define the nature of the behavioral deficits.

### Mutations of *nav2a* and *onecut1* do not impair the vestibulo-ocular reflex

*nav2a* is orthologous to *unc-53* in *C. elegans*, where it is essential for axonal and cell migration ^52^, and *sickie* in *D. melanogaster*, where loss-of-function is homozygous lethal ^53^. Mouse models of *Nav2* loss-of-function show cerebellar malformations, similar to a patient with a *NAV2* deletion who also exhibits deficits in voluntary eye movements ^53^. While the vestibulo-ocular reflex was normal in our *nav2a* mutants, their premature lethality is consistent with an important role for *nav2a* in development.

*onecut1* frameshift mutants had a normal vestibulo-ocular reflex and no obvious morphological defects. The *onecut* family is a highly conserved family of transcription factors whose members are implicated in development of various nervous system cell types ^54,55^, including upstream of *isl1* as regulators of spinal motor neuron diversity ^30^. Redundancy of these similar transcription factors ^56,57^ might compensate for the loss of *onecut1*. Compensation for mutations to *onecut1* may also occur through the process of transcriptional adaptation, in which degradation of mutant mRNA sequences triggers upregulation of similar genes ^58^. Mutagenesis targeting combinations of *onecut* factors (such as the zebrafish *onecut2, onecut3a, onecut3b*, and/or *onecutl*), or that creates an allele that fails to transcribe the mutant gene, thus eliminating the potential for degradation of mutant transcripts, could clarify *onecut* factors’ contributions to extraocular motor neuron development. Finally, though we did not observe any cellular, anatomical, or behavioral phenotypes through 7 dpf in homozygous mutant *onecut1* larvae, we cannot rule out the emergence of deficits later in development.

### Genes directly or indirectly implicated in oculomotor disorder

In addition to the genes selected for mutagenesis and subsequent experiments, our sequencing highlights several extraocular motor neuron subpopulation-specific genes that have been directly, or indirectly, implicated in oculomotor disorder. Duane Retraction Syndrome (DRS), an oculomotor disorder that presents with maldevelopment of the abducens nerve (nVI) and restricted horizontal eye movements, is caused by loss of *MAFB* function ^6,8^. This is consistent with recent analysis of gene expression in the zebrafish hindbrain showing transient *mafba* expression in hindbrain rhombomeres 5/6^59^, where the extraocular motor neurons that innervate the lateral rectus muscle aggregate in the abducens nucleus (nVI). In *Mafb* knockout mice, not only is nVI absent, but nIII shows aberrant branching as well ^6^. Interestingly, we observed *mafba* expression in trochlear (nIV) cells. This *mafba* expression was restricted to nIV, but only present at 50 hpf, at which point both dorsal nIII and nIV cells are in the process of extending their axons towards target muscles ^29^. Our transcriptomic and expression data expands the view of *MAFB* homologs in regulating extraocular muscle targeting.

Mutations in the gene *SEMA3F*, which encodes a member of the semaphorin signaling protein family, are associated with nIII hypoplasia ^8^. nIII axons defasciculate and fail to reach muscle partners in *sema3f* mutant zebrafish ^8^. Neuropilin 2 (Nrp2) is a receptor for Sema3f ^60^. Consistent with a role for Sema3f/Nrp2a signaling in oculomotor axon guidance, our *in situ* hybridization shows *nrp2a* RNA in early-born dorsal nIII cells. However, in bulk sequencing data, *nrp2a* was more strongly expressed in the late-born population . One possibility is that the expression is due to off-target binding of *nrp2a in situ* hybridization probes: the *in situ* expression pattern is similar to that of *nrp1a*, which was expressed most strongly in dorsal nIII at 50 hpf. While Neuropilin 1 (Nrp1) is a receptor for Sema3a ^61^, our data implicates class 3 semaphorins broadly as mediators of oculomotor axon guidance. Regulation of Nrp1 in zebrafish caudal primary (CaP) motor neurons contributes to CaP axon guidance in the spinal cord ^62^. Future work that similarly modulates levels of *nrp1a* expression in nIII cells would speak to the role of these guidance cues.

Trisomy 10 and duplications of the long arm of chromosome 10, which contains the intermediate filament encoding gene *INA*, have been linked to strabismus ^63,64^. Our *in situ* hybridization showed expression of the homolog *inab* specifically in nIV at 50, but not 33 or 80, hpf. By 50 hpf, nIV axons have crossed the midline and begun to grow ventrally towards the eye ^29^. Notably, *inab* is required for proper axon branching of subsets of zebrafish spinal motor neurons ^65^. Future investigations of extraocular motor neuron axonal morphology ^29^ could determine whether *inab* plays a similar role in extraocular motor neurons.

### Limitations

The major limitations of this study are common to all transcriptomic and loss-of-function work. While we used bulk sequencing, plate-based single-cell sequencing, and *in situ* validation, our datasets are unlikely to be comprehensive with respect to lowly-expressed genes. While it is beyond the scope of this paper, recent zebrafish datasets ^66–68^ include nIII/nIV; together with our data, these might provide a more complete picture of extraocular motor neuron transcriptomes. We only observed impaired vestibulo-ocular reflex behavior after loss of *sim1a*; as discussed above, we can only speculate with respect to why *nav2a* and *onecut1* did not show similar impairment.

More specifically, our reagents and choice of behavior limits what we can observe. The *Tg(isl1:GFP)* line we used to isolate extraocular motor neurons does not label inferior oblique or nVI (abducens) motor neurons ^15^. Future work with broader transgenic lines ^69^ would allow a more comprehensive characterization. Finally, our vestibulo-ocular reflex assay does not probe horizontal eye movements produced by the medial (nIII) or lateral rectus (nVI) motor neurons, and is limited with respect to stimulus kinetics. As discussed above, visual stimulation ^22^ and characterization of horizontal eye movements ^51^ could more comprehensively probe potential behavioral deficits.

## Conclusion

This study establishes a pipeline to discover the genes responsible for normal development of vertebrate extraocular motor neurons. Our assay reveals considerable diversity among developing motor neuron pools. Further, a loss-of-function approach coupled with a vestibuloocular reflex assay implicates *sim1a* in the development of extraocular motor neurons. Broadly, our work establishes a powerful approach to yield novel insights into the molecular mechanisms that govern development of the extraocular motor system in health and disease.

## MATERIALS AND METHODS

### Fish Care

All procedures involving zebrafish larvae (*Danio rerio*) were approved by the Institutional Animal Care and Use Committee of New York University Langone Health. Fertilized eggs were collected and maintained at 28.5° C on a standard 14/10 hour light/dark cycle. Before 5 dpf, larvae were maintained at densities of 20–50 larvae per petri dish of 10 cm diameter, filled with 25–40 mL E3 with 0.5 ppm methylene blue. After 5 dpf, larvae were maintained at densities under 20 larvae per petri dish and were fed cultured rotifers (Reed Mariculture) daily. Zebrafish larvae at the ages we studied have not yet differentiated their sex, and so sex was not considered as a behavioral variable.

### Transgenic Lines

The *Tg(isl1:Kaede)* line ^16^ was used for bulk sequencing experiments and *in situ* hybridization. The *Tg(isl1:GFP)* line ^15^ was used for single-cell sequencing, *in situ* hybridization, and motor neuron counts. Triple transgenic lines were used to identify the statoacoustic ganglion and tangential nucleus that carried the following alleles: *Tg(-17*.*6isl2b:GFP)* ^*70*^*;Tg(6*.*7Tru*.*Hcrtr2:GAL4-VP16)* ^*19,71*^; *Tg(UAS:E1b-Kaede)* ^*72*^. Motor neuron imaging experiments and *in situ* hybridization experiments were done on the *mitfa*^-/-^ background to remove pigment.

### Ocular Motor Neuron Birthdating

Kaede-expressing embryos were photoconverted as described in ^13^. Briefly, at 18, 22, 26, 30, 34, 42, 46, 50, or 54 hpf the entire fish was exposed to 405 nm LED light for 5 mins, and then raised in a dark incubator to prevent background photoconversion of Kaede fluorophore generated after the photoconversion timepoint. Photoconverted fish were then mounted dorsally and imaged at on a Zeiss LSM800 confocal microscope with a water-immersion objective (Zeiss W Plan-Apochromat 20x/1.0). To account for background conversion of Kaede due to incident light exposure during the experiment, non-converted controls were also imaged and were used to set a threshold level of red fluorescence. For each photoconversion timepoint, image stacks were manually evaluated in FIJI ^73^ to determine cell location and whether each cell was Kaede-red positive. Cells that were Kaede-red positive were considered born prior to that time-point.

### Ocular Motor Neuron Dissection, Dissociation, and Flow Cytometry

*Tg(isl1:Kaede)* larvae were photoconverted as above at 33 hpf by exposing the entire fish to a 405 nm LED light for 5 minutes. To prevent background photoconversion, larvae were raised in a dark incubator and were kept in the dark during motor neuron dissections. Ocular motor neurons were harvested from 50 hpf photoconverted larvae. Three experimenters (D.G., K.R.H., and P.L.) harvested neurons in parallel. Larvae were anesthetized in MESAB in Earle’s Balanced Salt Solution with calcium, magnesium, and phenol red (EBSS, Thermo Fisher Scientific 24010043) and larvae were positioned dorsal-up in an 3% agarose-molded petri dish with triangle wells. Fluorescence in motor neurons was visualized using a SugarCube LED Illuminator (Ushio America, Cypress CA) using 10x eyepieces on a stereomicroscope (Leica Microsystems, Wetzlar, Germany). Cells were harvested using a thin wall glass capillary tube (4 inch, OD 1.0MM, World Precision Instruments) into EBSS in a non-stick Eppendorf tube and kept on ice until dissociation. Cells were dissociated in 20 units/mL of papain prepared in EBSS (Worthington Biochemical), 2000 units/mL of deoxyribonuclease prepared in EBSS (Worthington Biochemical), and 100 mg/mL of Type 1A Collagenase (Sigma Aldrich) prepared in Hanks Buffered Salt Solution without calcium/magnesium (HBSS, Thermo Fisher Scientific). Cells were incubated for 45 minutes at 31.5° C with a gentle vortex every 10–15 min, then passed through a 20 µm filter and centrifuged for 10 mins at 300 x g. After removing supernatant, neurons were resuspended in L15 (Thermo Fisher Scientific) with 2% fetal bovine serum (Thermo Fisher Scientific). Cell health was evaluated using DAPI, applied at 0.5 µg/ml (Invitrogen) and incubated on ice for 30-45 mins prior to flow cytometry. Flow cytometry was performed using a Sony SH800z cell sorter (100 ţm nozzle, 20 psi) to isolate single neurons. Four controls were run for setting FACS gates: (1) non-fluorescent cells, (2) non-fluorescent cells + DAPI, (3) green fluorescent cells from unconverted *Tg(isl1:Kaede)* + DAPI and (4) red fluorescent cells from photoconverted *Tg(isl1:Kaede)* + DAPI. On average, less than 0.5% of cells were DAPI-positive and excluded. Experimental cells were gated for green fluorescence (late-born) or red + green fluorescence (early-born). Cells were sorted into an Eppendorf tube containing 700 µl of lysis buffer (RNAqueous Micro Total RNA Isolation Kit, Thermo Fisher Scientific) for downstream bulk RNA sequencing.

### Bulk RNA Sequencing and Analysis

RNA isolation was performed using an RNAqueous Micro Total RNA Isolation Kit (Thermo Fisher Scientific). RNA concentration and quality was evaluated using an RNA 6000 Pico Kit and a 2100 Bioanalyzer system (Agilent Technologies, Santa Clara, California). The majority of our samples had low RNA concentrations and accordingly did not have RIN scores, but for larger concentration samples RIN scores were routinely above 8 indicating that the extraction process did not affect RNA integrity. RNA sequencing was performed by NYU Langones Genome Technology Center (RRID: SCR_017929). Libraries were prepared using the low-input Clontech SMART-Seq HT with Nxt HTkit (Takara Bio USA) and sequenced using an Illumina NovaSeq 6000 with an S1 100Cycle Flow Cell (v1.5). Reads were mapped to GRCz11, and differential gene expression analysis was performed using the DESeq2 pipeline. Candidate genes were considered significantly differentially expressed if they had a fold-change greater than 2 and an adjusted p-value less than p=0.01.

Gene ontology (GO) analysis was performed in PANTHER19.0 (https://pantherdb.org/), testing for over-represented GO terms for molecular function among the 695 significantly differentially expressed genes ^74,75^. Statistical significance for GO terms was determined using a binomial test, with Bonferroni correction for multiple tests.

### Single-cell RNA Sequencing and Analysis

We dissected and dissociated nIII/nIV as above from 48 hpf *Tg(is1:GFP)* fish. We used a Sony SH800Z to FACS green cells and dispense them into plates for library preparation using a CEL-Seq2^28^ approach followed by sequencing performed by the NYU Genome Technology Center. After mapping reads to GRCz10, Seurat v4^76^ was used to exclude cells (<1,100 genes, >15,000 genes, >4% mitochondrial counts), normalize, and identify variable features. The top 2000 variable genes were used for dimensionality reduction by PCA; the top 21 principal components were retained for downstream analyses. Two 96-well plates yielded 84 cells that met our criteria, with 17,440±6,559 unique molecular identifiers per cell across 4,412±1,051 unique genes/cell.

### Fluorescent *In Situ* Hybridization and Imaging

Experiments were performed using Hybridization Chain Reaction (HCR) for whole-mount zebrafish larvae ^77^. Probes were generated using the HCR 3.0 probe maker using the sense sequence of the canonical gene cDNA from NCBI ^78^. Larvae were from the *Tg(isl1:Kaede)* background, photoconverted as above at 33 hpf. Pools of 6–8 larvae were fixed in 1.5 mL microcentrifuge tubes overnight with 4% PFA in 1x Phosphate-Buffered Saline (PBS) at 4°C and stored in 100% methanol at -20° C. Subsequently, HCR was performed as described in ^79^, with adjustments to proteinase K incubation time based on age (33 hpf: 17 minute incubation, 50 hpf: 21 minutes; 80 hpf: 33 minutes). HCR experiments used buffers and amplifiers from Molecular Instruments (Los Angeles, CA). Samples were stored in 1x PBS at 4° C until imaging. Larvae were mounted dorsally in 2% low-melting temperature agarose (Invitrogen 16520-050) in 1x PBS and imaged on a Zeiss LSM800 confocal microscope with 20x water-dipping objective. Laser power was kept consistent for all fluorophores across all imaged fish. White levels in representative images have been been adjusted for optimal viewing of fluorescence signal; all images from the same channel within the same figure have been adjusted to have the same minimum and maximum pixel intensity to facilitate comparison within figures.

### CRISPR-Cas9 Mutagenesis

Gene-specific CRISPR target sequences were selected using CRISPRscan ^80^, and were synthesized as DNA oligos with a 5’ T7 promoter (TAATACGACTCACTATA) and a 3’ overlap region (GTTTTAGAGCTAGAA). The DNA target sequence for *nav2a* was AGCGCCGCCCCGGTGGCCAA, for *sim1a* was GGAGGGCAGAGGCAGCAGTT, and for *onecut1* was GTGGCCCCGGGGTGCGCGTACGG. Single guide RNA (sgRNA) was generated as described in ^81^, with the following modifications: *in vitro* transcription was performed using the AmpliScribe T7 Flash Transcription Kit (LGC Biosearch Technologies) according to the manufacturer protocol and allowing the transcription reaction to incubate at 37° for 4 hours. sgRNA was precipitated using sodium acetate as described in ^82^. Crispants were generated by injecting zebrafish embryos at the 1-cell stage with 1 nL of injection mixture containing a final concentration of 4 uM EnGen Spy Cas9 NLS protein (NEB M0646), 100 ng/µl sgRNA, and 10% phenol red in a Cas9 protein buffer solution (10 mM MgCl_2_, 200 mM KCl, 20 mM Tris Buffer pH 8.0, ^81^). Injection mixture was incubated at 37°C for 5 minutes prior to injections to promote formation of the Cas9/sgRNA complex.

Injected fish were then raised and founders (F0) identified by out-crossing and sequencing pools of 10–20 embryos using Sanger sequencing of an approximately 500 base pair region around the target sequence. One allele of each gene was identified and used for experiments: *sim1a*^*d17*^ has a 17 bp deletion from base pairs 693 to 709. *nav2a*^*d8*^ has an 8 bp deletion from base pairs 151590 to 151597. *onecut1*^*d14*^ has a 14 bp deletion from base pairs 826 to 839. Each mutation was a frameshift mutation, causing a predicted premature stop codon. Studies using *phox2a, nav2a*, and *onecut1* stable line mutants were performed on larvae from in-crosses of F2 or later generations. For *sim1a*, studies were performed on larvae from in-crosses of the F1 generation.

### Eye Rotation Behavioral Assay

Body tilts were delivered and eye rotations measured and analyzed as per ^21^. Briefly, 6–7 dpf larvae were mounted in 2% low-melting temperature agarose on a piece of Sylgard 184 (Dow Corning), the left eye was freed, and the Sylgard was placed in a 10 mm optical glass cuvette filled with 1 mL of E3. The cuvette was then mounted to a rotating platform. The eye was imaged with a 5x objective (Olympus MPLN, 0.1 NA) onto a machine vision camera (Guppy Pro 2 F-031, Allied Vision Technologies) which acquired a 100x100 pixel image of the left eye of the fish (6 µm/pixel) at 200 Hz. Images were processed on-line to derive an estimate of torsional angle (LabView 2014, National Instruments), and data was analyzed using custom MATLAB scripts (Mathworks, Natick MA).

Each experiment consisted of 50 cycles of four steps. Steps were ± 15° towards and away from the horizon, with a trapezoidal velocity profile that peaked at 35°/sec, peak acceleration 150°/sec^2^. Eye and platform data was processed as per ^21^; each step was evaluated manually to exclude trials with rapid deviations in eye position indicative of horizontal saccades or gross failure of the pattern-matching algorithm. The response to each step for a given fish was defined as the peak eye velocity occurring over the first second after the start of the step, averaged across all cycles.

### Ocular motor neuron imaging and counts

Larvae between 5–7 dpf were mounted dorsally in 2% low-melting temperature agarose in E3 and imaged on a Zeiss LSM800 confocal microscope with a water-immersion objective (Zeiss W Plan-Apochromat 20x/1.0). Stacks spanned ∼90 µM, sampled every 1.5 µM. Analysis was performed in Fiji/ImageJ using the Cell Counter plugin. Stacks of nIII/nIV were subdivided in the dorsoventral axis as described in ^14^ to facilitate localization. A point ROI was dropped over each neuron in the plane in which the soma was brightest, and the number of neurons belonging to nIV, dorsal nIII, and ventral nIII in each plane was recorded.

### Data & Code

All data, raw and analyzed, as well as code necessary to generate the figures will be available at 10.17605/OSF.IO/JR4GS

## ACKNOWLEDGMENTS

Research was supported by the National Eye Institute under award number R01EY035691, the National Institute on Deafness and Communication Disorders of the National Institutes of Health under award numbers R01DC017489, F31DC019554, and F31DC020910, and the National Institute of Neurological Disorders and Stroke under award numbers F99NS129179, T32NS086750, and the National Cancer Institute P30CA016087. The authors would like to thank Elizabeth Engle for confirming the absence of known SIM1 human variants, Itai Yanai and Felicia Kuperwaser for help with CEL-Seq2, the NYU Langone Health Genome Technology Center for sequencing, alignment, and analysis, and members of the Schoppik and Nagel Labs for their helpful feedback.

## AUTHOR CONTRIBUTIONS

Conceptualization: DS, KRH; Methodology: KRH, DG, BR; Investigation: KRH, BR, HG, GX, CQ, DG, PL, EG; Visualization: KRH, EG; Writing: EG, KRH; Editing: DS, EG; Funding Acquisition: KRH, DS; Supervision: KRH, DS.

---

**Figure S1:**
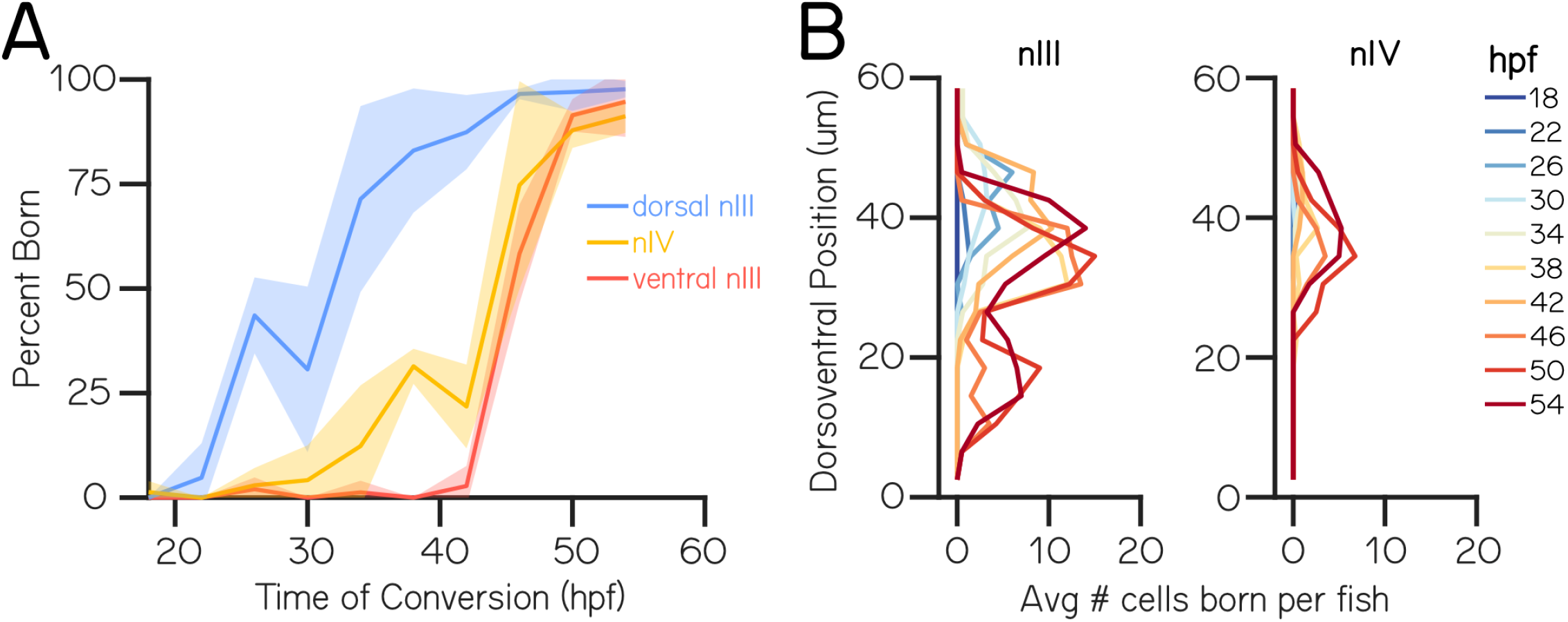
extraocular motor neuron birthdate varies with dorsoventral position. (A) Percent of *Tg(isl1:Kaede)* neurons that were born prior to the time of photoconversion in dorsal nIII (blue), ventral nIII (red) and nIV (yellow). Lines are averages across all fish, ribbons represent *±* 1 standard deviation or range. N = 4 fish (18 hpf), 3 fish (22 hpf), 2 fish (26 hpf), 4 fish (30 hpf), 5 fish (34 hpf), 3 fish (42 hpf), 2 fish (46 hpf), 4 fish (50 hpf), and 4 fish (54 hpf) (B)Dorsoventral position of photoconverted *Tg(isl1:Kaede)* neurons in nIII (left) and nIV (right) colored by photoconversion timepoint (18-54 hpf).

**Figure S2:**
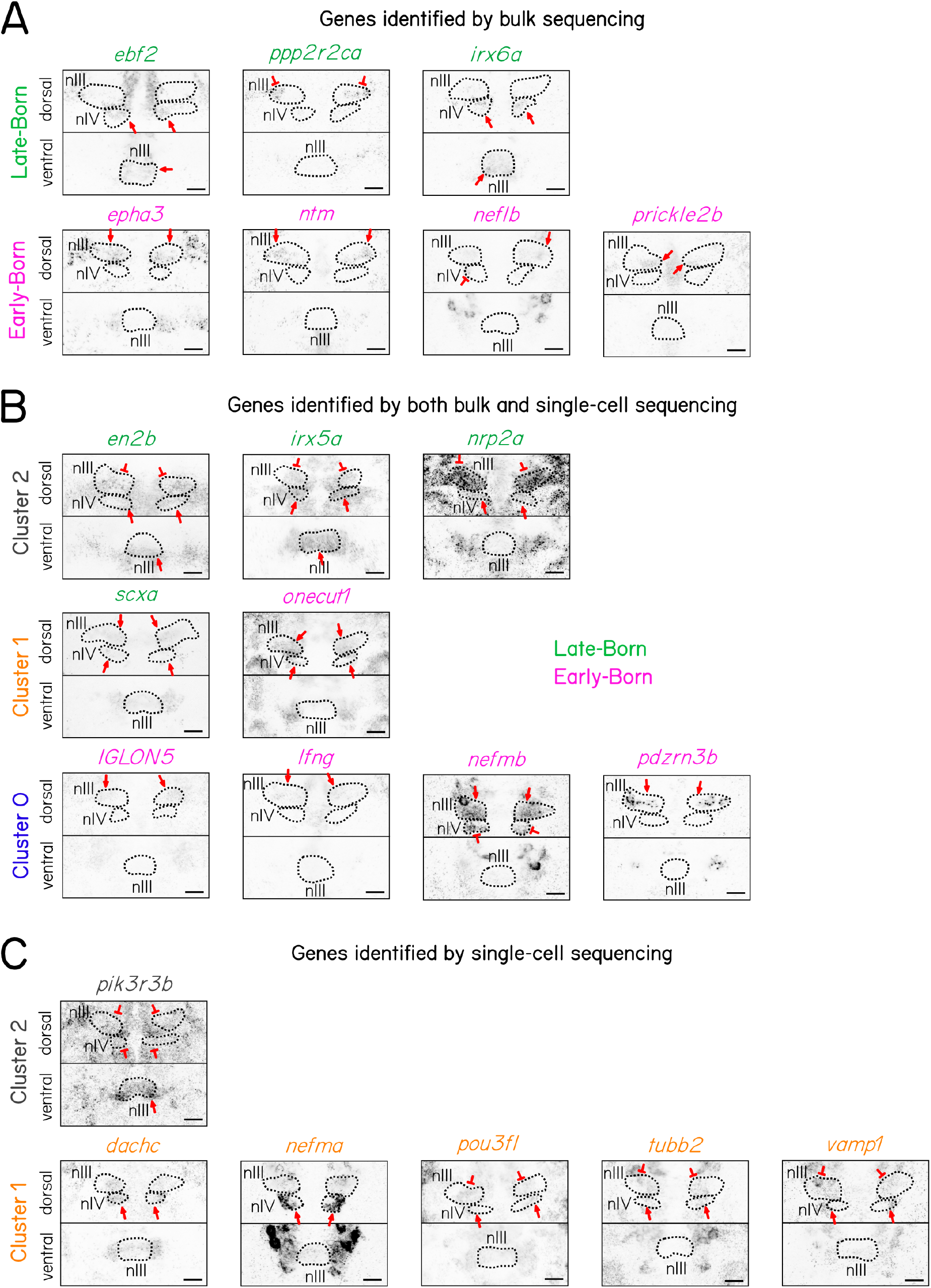
Fluorescent *in situ* hybridization expression for candidate genes identified in bulk and single-cell RNA sequencing of extraocular motor neurons in nIII and nIV. Maximum intensity projections of RNA probe expression in extraocular motor neurons split by dorsal (dorsal nIII and nIV) and ventral (ventral nIII) location. Black dashed outlines correspond to the location of extraocular motor neuron populations labeled by *Tg(isl1:Kaede)* (Kaede expression not shown). Scale bar 20 µm. **(A)** RNA probe expression for genes identified by bulk sequencing of late- (top, green) and early-born (bottom, pink) cells. Red arrows indicate expression in populations in which candidates were predicted to be enriched based on sequencing data. Red Ts indicate expression in other populations. **(B)** RNA probe expression for genes identified by both bulk and single-cell sequencing. Gene names are colored by late- (green) and early-born (pink) groups and organized by cluster (top: cluster 0, gray; center: cluster 1, orange; bottom: cluster 2, blue). Red arrows indicate expression in populations in which candidates were predicted to be enriched based on sequencing data. Red Ts indicate expression in other populations. **(C)** RNA probe expression for genes identified by single-cell sequencing. Genes are organized by cluster (top: cluster 0, gray; bottom: cluster 1, orange). Red arrows indicate expression in populations in which candidates were predicted to be enriched based on sequencing data. Red Ts indicate expression in other populations.

**Figure S3:**
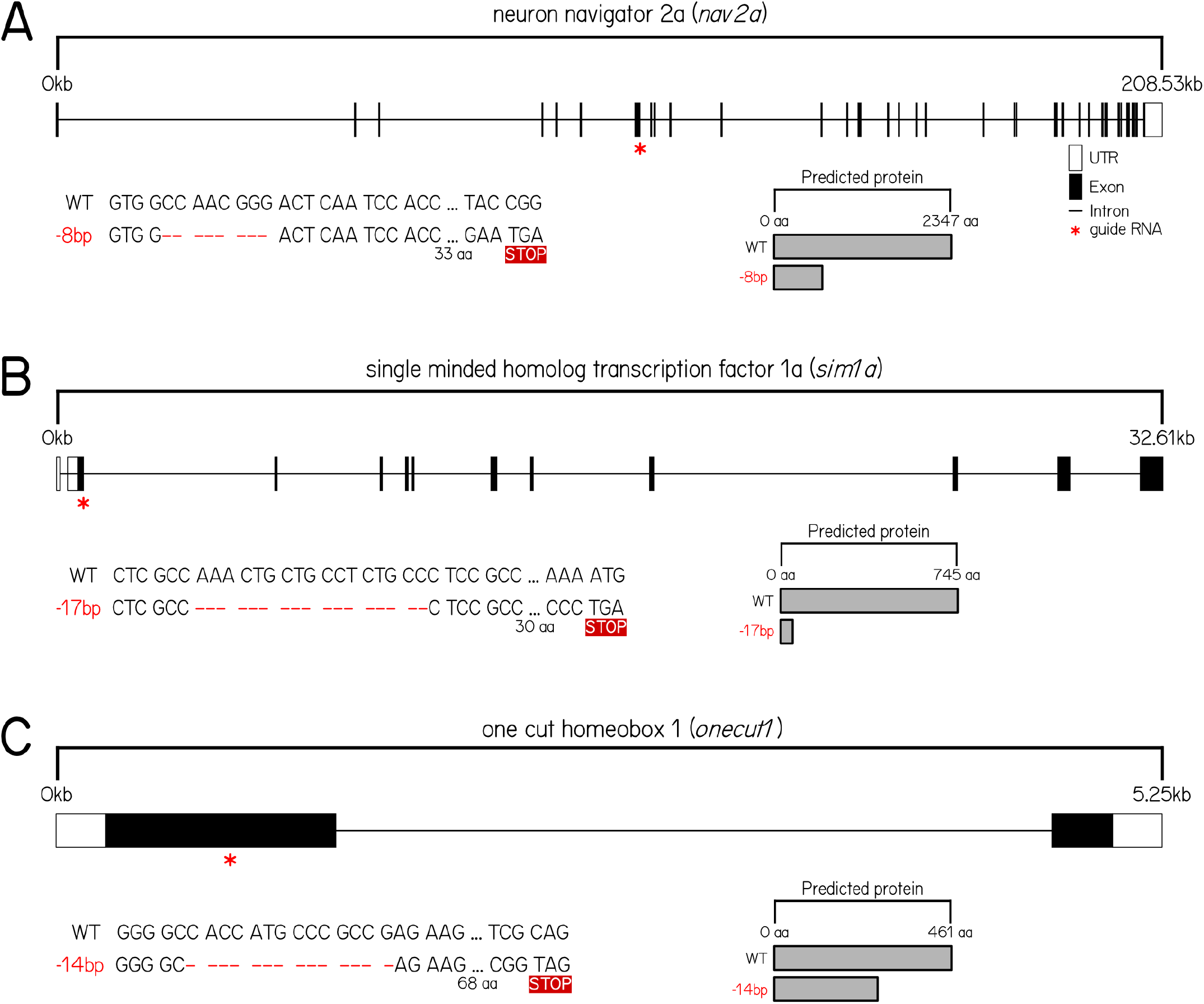
CRISPR/Cas9 mediated mutagenesis of select candidate genes. **(A)** Top: Schematic representing mutation created in *nav2a*. Red star indicates location of guides against nav2a DNA. Filled boxes show exons, open boxes show UTRs, and horizontal lines show introns. Bottom: Left shows DNA sequence in wildtype and *nav2a*^*d8*^ alleles. Red dashed lines indicate deleted sequence. STOP box indicates predicted premature stop codon due to deletion. Right shows predicted protein sequence. **(B)** Top: Schematic representing mutation created in *sim1a*. Bottom: DNA sequence and predicted protein in wildtype and *sim1a*^*d17*^ alleles. **(C)** Top: Schematic representing mutation created in *onecut1*. Bottom: DNA sequence and predicted protein in wildtype and *onecut1*^*d14*^ alleles.

## REFERENCES

[1] Linda R. Dagi, Yoon-Hee Chang, and Evan Silverstein. Complex or Incomitant Strabismus, page 122. Springer International Publishing, 2020.

[2] Eileen E. Birch. Amblyopia and binocular vision. Progress in Retinal and Eye Research, 33:6784, March 2013.

[3] Sarah R. Hatt, David A. Leske, Yolanda S. Castañeda, Suzanne M. Wernimont, Laura Liebermann, Christina S. Cheng-Patel, Eileen E. Birch, and Jonathan M. Holmes. Association of strabismus with functional vision and eye-related quality of life in children. JAMA Ophthalmology, 138(5):528, May 2020.

[4] Mary C. Whitman and Elizabeth C. Engle. Ocular congenital cranial dysinnervation disorders (ccdds): insights into axon growth and guidance. Human Molecular Genetics, 26(R1):R37R44, April 2017.

[5] Mary C. Whitman and Elizabeth C. Engle. Genetics of Strabismus, page 68876905. Springer International Publishing, 2022.

[6] Jong G. Park, Max A. Tischfield, Alicia A. Nugent, Long Cheng, Silvio Alessandro Di Gioia, Wai-Man Chan, Gail Maconachie, Thomas M. Bosley, C. Gail Summers, David G. Hunter, Caroline D. Robson, Irene Gottlob, and Elizabeth C. Engle. Loss of mafb function in humans and mice causes duane syndrome, aberrant extraocular muscle innervation, and inner-ear defects. The American Journal of Human Genetics, 98(6):12201227, June 2016.

[7] Motoi Nakano, Koki Yamada, Jennifer Fain, Emin C. Sener, Carol J. Selleck, Abdulaziz H. Awad, Johan Zwaan, Paul B. Mullaney, Thomas M. Bosley, and Elizabeth C. Engle. Homozygous mutations in arix(phox2a) result in congenital fibrosis of the extraocular muscles type 2. Nature Genetics, 29(3):315320, October 2001.

[8] Julie A. Jurgens, Paola M. Matos Ruiz, Jessica King, Emma E. Foster, Lindsay Berube, Wai-Man Chan, Brenda J. Barry, Raehoon Jeong, Elisabeth Rothman, Mary C. Whitman, Sarah MacKinnon, Cristina Rivera-Quiles, Brandon M. Pratt, Teresa Easterbrooks, Fiona M. Mensching, Silvio Alessandro Di Gioia, Lynn Pais, Eleina M. England, Teresa de Berardinis, Adriano Magli, Feray Koc, Kazuhide Asakawa, Koichi Kawakami, Anne O’Donnell-Luria, David G. Hunter, Caroline D. Robson, Martha L. Bulyk, and Elizabeth C. Engle. Gene identification for ocular congenital cranial motor neuron disorders using human sequencing, zebrafish screening, and protein binding microarrays. Investigative Ophthalmology & Visual Science, (62), March 2025.

[9] Sarah Guthrie. Patterning and axon guidance of cranial motor neurons. Nature Reviews Neuroscience, 8(11):859871, November 2007.

[10] Robert F. Spencer and John D. Porter. Biological organization of the extraocular muscles, page 4380. Elsevier, 2006.

[11] J.A. Büttner-Ennever. The extraocular motor nuclei: organization and functional neuroanatomy, page 95125. Elsevier, 2006.

[12] Dena Goldblatt, Stephanie Huang, Marie R. Greaney, Kyla R. Hamling, Venkatakaushik Voleti, Citlali Perez-Campos, Kripa B. Patel, Wenze Li, Elizabeth M.C. Hillman, Martha W. Bagnall, and David Schoppik. Neuronal birthdate reveals topography in a vestibular brainstem circuit for gaze stabilization. Current Biology, 33(7):1265–1281.e7, April 2023.

[13] Stephanie Huang, Emily Gershowitz, Marie R. Greaney, Samantha N. Davis, David Schoppik, and Dena Goldblatt. Birthdate aligns vestibular sensory neurons with central and motor partners across a sensorimotor reflex circuit for gaze stabilization. Development, January 2026.

[14] Marie R. Greaney, Ann E. Privorotskiy, Kristen P. D’Elia, and David Schoppik. Extraocular motoneuron pools develop along a dorsoventral axis in zebrafish, Danio rerio. Journal of Comparative Neurology, 525(1):65–78, June 2016.

[15] Shin-ichi Higashijima, Yoshiki Hotta, and Hitoshi Okamoto. Visualization of cranial motor neurons in live transgenic zebrafish expressing green fluorescent protein under the control of theislet-1promoter/enhancer. The Journal of Neuroscience, 20(1):206218, January 2000.

[16] Gabrielle R. Barsh, Adam J. Isabella, and Cecilia B. Moens. Vagus motor neuron topographic map determined by parallel mechanisms of hox5 expression and time of axon initiation. Current Biology, 27(24):3812–3825.e3, December 2017.

[17] Celine Bellegarda, Franziska Auer, and David Schoppik. Zebrafish as a model to understand extraocular motor neuron diversity. Current Opinion in Neurobiology, 90:102964, February 2025.

[18] Isaac H. Bianco, Leung-Hang Ma, David Schoppik, Drew N. Robson, Michael B. Orger, James C. Beck, Jennifer M. Li, Alexander F. Schier, Florian Engert, and Robert Baker. The tangential nucleus controls a gravito-inertial vestibulo-ocular reflex. Current Biology, 22(14):12851295, July 2012.

[19] David Schoppik, Isaac H. Bianco, David A. Prober, Adam D. Douglass, Drew N. Robson, Jennifer M.B. Li, Joel S.F. Greenwood, Edward Soucy, Florian Engert, and Alexander F. Schier. Gaze-stabilizing central vestibular neurons project asymmetrically to extraocular motoneuron pools. The Journal of Neuroscience, 37(47):1135311365, September 2017.

[20] Dena Goldblatt, Baak Rosti, Kyla R Hamling, Paige Leary, Harsh Panchal, Marlyn Li, Hannah Gelnaw, Stephanie Huang, Cheryl Quainoo, and David Schoppik. Motor neurons are dispensable for the assembly of a sensorimotor circuit for gaze stabilization. eLife, October 2024.

[21] Paige Leary, Celine Bellegarda, Cheryl Quainoo, Dena Goldblatt, Baak Rosti, and David Schoppik. Sensation is dispensable for the maturation of the vestibulo-ocular reflex. Science, 387(6729):8590, January 2025.

[22] David-Samuel Burkhardt, Gabriel Möller, Laurian Deligand, Christiane Fichtner, Tim C. Hladnik, and Aristides B. Arrenberg. Vertical optokinetic eye movements in the larval zebrafish. bioRxiv, March 2025.

[23] Zhikai Liu, David G. C. Hildebrand, Joshua L. Morgan, Yizhen Jia, Nicholas Slimmon, and Martha W. Bagnall. Organization of the gravity-sensing system in zebrafish. Nature Communications, 13(1), August 2022.

[24] Kristen P. D’Elia, Hanna Hameedy, Dena Goldblatt, Paul Frazel, Mercer Kriese, Yunlu Zhu, Kyla R. Hamling, Koichi Kawakami, Shane A. Liddelow, David Schoppik, and Jeremy S. Dasen. Determinants of motor neuron functional subtypes important for locomotor speed. Cell Reports, 42(9):113049, September 2023.

[25] Irene Pallucchi, Maria Bertuzzi, David Madrid, Pierre Fontanel, Shin-ichi Higashijima, and Abdeljabbar El Manira. Molecular blueprints for spinal circuit modules controlling locomotor speed in zebrafish. Nature Neuroscience, 27(1):7889, November 2023.

[26] Jimmy J Kelly, Hua Wen, and Paul Brehm. Single-cell rnaseq analysis of spinal locomotor circuitry in larval zebrafish. eLife, 12, November 2023.

[27] Craig Evinger. Extraocular motor nuclei: location, morphology and afferents. Reviews of Oculomotor Research, 2:81–117, 1988.

[28] Tamar Hashimshony, Naftalie Senderovich, Gal Avital, Agnes Klochendler, Yaron de Leeuw, Leon Anavy, Dave Gennert, Shuqiang Li, Kenneth J. Livak, Orit Rozenblatt-Rosen, Yuval Dor, Aviv Regev, and Itai Yanai. CEL-seq2: sensitive highly-multiplexed single-cell RNA-seq. Genome Biology, 17(1), April 2016.

[29] Christopher Clark, Oliver Austen, Ivana Poparic, and Sarah Guthrie. 2-chimaerin regulates a key axon guidance transition during development of the oculomotor projection. The Journal of Neuroscience, 33(42):1654016551, October 2013.

[30] Agnès Roy, Cédric Francius, David L. Rousso, Eve Seuntjens, Joke Debruyn, Georg Luxenhofer, Andrea B. Huber, Danny Huylebroeck, Bennett G. Novitch, and Frédéric Clotman. Onecut transcription factors act upstream of isl1 to regulate spinal motoneuron diversification. Development, 139(17):31093119, September 2012.

[31] Stephen T Crews and Chen-Ming Fan. Remembrance of things pas: regulation of development by bhlhpas proteins. Current Opinion in Genetics & Development, 9(5):580587, October 1999.

[32] Stephen T. Crews, John B. Thomas, and Corey S. Goodman. The drosophila single-minded gene encodes a nuclear protein with sequence similarity to the per gene product. Cell, 52(1):143151, January 1988.

[33] Chen-Ming Fan, Ellen Kuwana, Alessandro Bulfone, Colin F. Fletcher, Neal G. Copeland, Nancy A. Jenkins, Stephen Crews, Salvador Martinez, Luis Puelles, John L.R. Rubenstein, and Marc Tessier-Lavigne. Expression patterns of two murine homologs ofdrosophila single-mindedsuggest possible roles in embryonic patterning and in the pathogenesis of down syndrome. Molecular and Cellular Neuroscience, 7(1):116, January 1996.

[34] Roman Chrast, Hamish S. Scott, Haiming Chen, Jun Kudoh, Colette Rossier, Shinsei Minoshima, Yimin Wang, Nobuyoshi Shimizu, and Stylianos E. Antonarakis. Cloning of two human homologs of the drosophila single-minded gene sim1 on chromosome 6q and sim2 on 21q within the down syndrome chromosomalregion. Genome Research, 7(6):615624, June 1997.

[35] Fabrizio C. Serluca and Mark C. Fishman. Pre-pattern in the pronephric kidney field of zebrafish. Development, 128(12):22332241, June 2001.

[36] Christina N. Cheng and Rebecca A. Wingert. Nephron proximal tubule patterning and corpuscles of stannius formation are regulated by the sim1a transcription factor and retinoic acid in zebrafish. Developmental Biology, 399(1):100116, March 2015.

[37] Bridgette E. Drummond, Yue Li, Amanda N. Marra, Christina N. Cheng, and Rebecca A. Wingert. The tbx2a/b transcription factors direct pronephros segmentation and corpuscle of stannius formation in zebrafish. Developmental Biology, 421(1):5266, January 2017.

[38] Nataliya Borodovsky, Tatyana Ponomaryov, Shani Frenkel, and Gil Levkowitz. Neural protein olig2 acts upstream of the transcriptional regulator sim1 to specify diencephalic dopaminergic neurons. Developmental Dynamics, 238(4):826834, March 2009.

[39] Jennifer L. Eaton and Eric Glasgow. The zebrafish bhlh pas transcriptional regulator, singleminded 1 (sim1), is required for isotocin cell development. Developmental Dynamics, 235(8):20712082, July 2006.

[40] Heiko Lohr, Soojin Ryu, and Wolfgang Driever. Zebrafish diencephalic a11-related dopaminergic neurons share a conserved transcriptional network with neuroendocrine cell lineages. Development, 136(6):10071017, March 2009.

[41] Andrea Wolf and Soojin Ryu. Specification of posterior hypothalamic neurons requires coordinated activities of fezf2, otp, sim1a and foxb1.2. Development, 140(8):17621773, April 2013.

[42] Craig T. Jacobs, Aarti Kejriwal, Katrinka M. Kocha, Kevin Y. Jin, and Peng Huang. Temporal cell fate determination in the spinal cord is mediated by the duration of notch signalling. Developmental Biology, 489:113, September 2022.

[43] Jörn Schweitzer, Heiko Löhr, Joshua L. Bonkowsky, Katrin Hübscher, and Wolfgang Driever. Sim1a and arnt2 contribute to hypothalamo-spinal axon guidance by regulating robo2 activity via a robo3-dependent mechanism. Development, 140(1):93106, January 2013.

[44] Dylan Deska-Gauthier, Joanna Borowska-Fielding, Christopher T. Jones, and Ying Zhang. The temporal neurogenesis patterning of spinal p3v3 interneurons into divergent sub-population assemblies. The Journal of Neuroscience, 40(7):14401452, December 2019.

[45] Kyla R. Hamling, Yunlu Zhu, Franziska Auer, and David Schoppik. Tilt in place microscopy: a simple, low-cost solution to image neural responses to body rotations. The Journal of Neuroscience, 43(6):936948, December 2022.

[46] Monica C. Varela, Alex Y. Simões-Sato, Chong A. Kim, Débora R. Bertola, Claudia I.E. De Castro, and Celia P. Koiffmann. A new case ofăinterstitial 6q16.2ădeletion inăaăpatient with praderwilli-like phenotype andăinvestigation ofăsim1 gene deletion ină87ăpatients with syndromic obesity. European Journal of Medical Genetics, 49(4):298305, July 2006.

[47] Khaled K. Abu-Amero, Ali Hellani, Mustafa A. Salih, Abdulkarim Al Hussain, Majed al Obailan, Ghassan Zidan, Ibrahim A. Alorainy, and Thomas M. Bosley. Ophthalmologic abnormalities in a de novo terminal 6q deletion. Ophthalmic Genetics, 31(1):111, February 2010.

[48] Estephania Candelo, Max M. Feinstein, Diana Ramirez-Montaño, Juan F. Gomez, and Harry Pachajoa. First case report of praderwilli-like syndrome in colombia. Frontiers in Genetics, 9, March 2018.

[49] Tetsuya Okazaki, Tatsuya Kawaguchi, Yusuke Saiki, Chisako Aoki, Noriko Kasagi, Kaori Adachi, Ken Saida, Naomichi Matsumoto, Eiji Nanba, and Yoshihiro Maegaki. Clinical course of a japanese patient with developmental delay linked to a small 6q16.1 deletion. Human Genome Variation, 9(1), May 2022.

[50] Agnes M.F Wong, James A Sharpe, and Douglas Tweed. The vestibulo-ocular reflex in fourth nerve palsy: deficits and adaptation. Vision Research, 42(18):22052218, August 2002.

[51] Charles K. Dowell, Thomas Hawkins, and Isaac H. Bianco. Subsets of extraocular motoneurons produce kinematically distinct saccades during hunting and exploration. Current Biology, 35(3):554–573.e6, February 2025.

[52] Eve Stringham, Nathalie Pujol, Joel Vandekerckhove, and Thierry Bogaert. unc-53controls longitudinal migration inc. elegans. Development, 129(14):33673379, July 2002.

[53] Andrea Accogli, Shenzhao Lu, Ilaria Musante, Paolo Scudieri, Jill A. Rosenfeld, Mariasavina Severino, Simona Baldassari, Michele Iacomino, Antonella Riva, Ganna Balagura, Gianluca Piccolo, Carlo Minetti, Denis Roberto, Fan Xia, Razaali Razak, Emily Lawrence, Mohamed Hussein, Emmanuel Yih-Herng Chang, Michelle Holick, Elisa Calì, Emanuela Aliberto, Rosalba De-Sarro, Antonio Gambardella, Undiagnosed Diseases Network, SYNaPS Study Group, Lisa Emrick, Peter J. A. McCaffery, Margaret Clagett-Dame, Paul C. Marcogliese, Hugo J. Bellen, Seema R. Lalani, Federico Zara, Pasquale Striano, and Vincenzo Salpietro. Loss of neuron navigator 2 impairs brain and cerebellar development. The Cerebellum, 22(2):206222, February 2022.

[54] Andreas Sagner, Isabel Zhang, Thomas Watson, Jorge Lazaro, Manuela Melchionda, and James Briscoe. A shared transcriptional code orchestrates temporal patterning of the central nervous system. PLOS Biology, 19(11):e3001450, November 2021.

[55] Ho Sung Rhee, Michael Closser, Yuchun Guo, Elizaveta V. Bashkirova, G. Christopher Tan, David K. Gifford, and Hynek Wichterle. Expression of terminal effector genes in mammalian neurons is maintained by a dynamic relay of transient enhancers. Neuron, 92(6):12521265, December 2016.

[56] Vinciane Vanhorenbeeck, Marjorie Jenny, Jean-François Cornut, Gérard Gradwohl, Frédéric P. Lemaigre, Guy G. Rousseau, and Patrick Jacquemin. Role of the onecut transcription factors in pancreas morphogenesis and in pancreatic and enteric endocrine differentiation. Developmental Biology, 305(2):685694, May 2007.

[57] Quirino Attilio Vassalli, Giulia Fasano, Valeria Nittoli, Eleonora Gagliardi, Rosa Maria Sepe, Aldo Donizetti, Francesco Aniello, Paolo Sordino, Robert Kelsh, and Annamaria Locascio. The zebrafish retina and the evolution of the onecut-mediated pathway in cell type differentiation. Cells, 13(24):2071, December 2024.

[58] Mohamed A. El-Brolosy, Zacharias Kontarakis, Andrea Rossi, Carsten Kuenne, Stefan Günther, Nana Fukuda, Khrievono Kikhi, Giulia L. M. Boezio, Carter M. Takacs, Shih-Lei Lai, Ryuichi Fukuda, Claudia Gerri, Antonio J. Giraldez, and Didier Y. R. Stainier. Genetic compensation triggered by mutant mrna degradation. Nature, 568(7751):193197, April 2019.

[59] Monica Tambalo, Richard Mitter, and David G. Wilkinson. A single cell transcriptome atlas of the developing zebrafish hindbrain. Development, 147(6), March 2020.

[60] Andrea B. Huber, Artur Kania, Tracy S. Tran, Chenghua Gu, Natalia De Marco Garcia, Ivo Lieberam, Dontais Johnson, Thomas M. Jessell, David D. Ginty, and Alex L. Kolodkin. Distinct roles for secreted semaphorin signaling in spinal motor axon guidance. Neuron, 48(6):949964, December 2005.

[61] Hajime Fujisawa and Takashi Kitsukawa. Receptors for collapsin/semaphorins. Current Opinion in Neurobiology, 8(5):587592, October 1998.

[62] Mika Sato-Maeda, Hiroshi Tawarayama, Masuo Obinata, John Y. Kuwada, and Wataru Shoji. Sema3a1 guides spinal motor axons in a cell- and stage-specific manner in zebrafish. Development, 133(5):937947, March 2006.

[63] Catherine Boon, Tom Wlarkello, Colleen JacksonCook, and Arti Pandya. Partial trisomy 10 mosaicism with cutaneous manifestations: report of a case and review of the literature. Clinical Genetics, 50(5):417421, November 1996.

[64] J. P. Fryns, A. Kleczkowska, L. IgodtAmeye, and H. Van den Berghe. Proximal duplication of the long arm of chromosome 10 (10q11.2 10q22): a distinct clinical entity. Clinical Genetics, 32(1):6165, July 1987.

[65] Liesl Van Ryswyk, Levi Simonson, and Judith S. Eisen. The role of inab in axon morphology of an identified zebrafish motoneuron. PLoS ONE, 9(2):e88631, February 2014.

[66] Dylan R. Farnsworth, Lauren M. Saunders, and Adam C. Miller. A single-cell transcriptome atlas for zebrafish development. Developmental Biology, 459(2):100108, March 2020.

[67] Lauren M. Saunders, Sanjay R. Srivatsan, Madeleine Duran, Michael W. Dorrity, Brent Ewing, Tor H. Linbo, Jay Shendure, David W. Raible, Cecilia B. Moens, David Kimelman, and Cole Trapnell. Embryo-scale reverse genetics at single-cell resolution. Nature, 623(7988):782791, November 2023.

[68] Lior Fishman, Gal Nechooshtan, Florian Erhard, Aviv Regev, Jeffrey A. Farrell, and Michal Rabani. Single-cell temporal dynamics reveals the relative contributions of transcription and degradation to cell-type specific gene expression in zebrafish embryos. bioRxiv, April 2023.

[69] Atsushi Taniguchi, Yukiko Kimura, Ikue Mori, Shigenori Nonaka, and Shinichi Higashijima. Axiallyconfined in vivo singlecell labeling by primed conversion using blue and red lasers with conventional confocal microscopes. Development, Growth & Differentiation, 59(9):741748, December 2017.

[70] A. J. Pittman, M.-Y. Law, and C.-B. Chien. Pathfinding in a large vertebrate axon tract: isotypic interactions guide retinotectal axons at multiple choice points. Development, 135(17):2865–2871, sep 2008.

[71] AlixăM.B. Lacoste, David Schoppik, DrewăN. Robson, Martin Haesemeyer, Ruben Portugues, JenniferăM. Li, Owen Randlett, CarolineăL. Wee, Florian Engert, and AlexanderăF. Schier. A convergent and essential interneuron pathway for mauthner-cell-mediated escapes. Current Biology, 25(11):15261534, June 2015.

[72] Ethan K Scott, Lindsay Mason, Aristides B Arrenberg, Limor Ziv, Nathan J Gosse, Tong Xiao, Neil C Chi, Kazuhide Asakawa, Koichi Kawakami, and Herwig Baier. Targeting neural circuitry in zebrafish using gal4 enhancer trapping. Nature Methods, 4(4):323326, March 2007.

[73] Johannes Schindelin, Ignacio Arganda-Carreras, Erwin Frise, Verena Kaynig, Mark Longair, Tobias Pietzsch, Stephan Preibisch, Curtis Rueden, Stephan Saalfeld, Benjamin Schmid, Jean-Yves Tinevez, Daniel James White, Volker Hartenstein, Kevin Eliceiri, Pavel Tomancak, and Albert Cardona. Fiji: an open-source platform for biological-image analysis. Nature Methods, 9(7):676–682, June 2012.

[74] Paul D. Thomas, Dustin Ebert, Anushya Muruganujan, Tremayne Mushayahama, LaurentPhilippe Albou, and Huaiyu Mi. <scp>panther</scp>: Making genomescale phylogenetics accessible to all. Protein Science, 31(1):822, November 2021.

[75] Huaiyu Mi, Anushya Muruganujan, Xiaosong Huang, Dustin Ebert, Caitlin Mills, Xinyu Guo, and Paul D. Thomas. Protocol update for large-scale genome and gene function analysis with the panther classification system (v.14.0). Nature Protocols, 14(3):703721, February 2019.

[76] Yuhan Hao, Stephanie Hao, Erica Andersen-Nissen, William M. Mauck, Shiwei Zheng, Andrew Butler, Maddie J. Lee, Aaron J. Wilk, Charlotte Darby, Michael Zager, Paul Hoffman, Marlon Stoeckius, Efthymia Papalexi, Eleni P. Mimitou, Jaison Jain, Avi Srivastava, Tim Stuart, Lamar M. Fleming, Bertrand Yeung, Angela J. Rogers, Juliana M. McElrath, Catherine A. Blish, Raphael Gottardo, Peter Smibert, and Rahul Satija. Integrated analysis of multimodal single-cell data. Cell, 184(13):3573–3587.e29, June 2021.

[77] Harry M. T. Choi, Maayan Schwarzkopf, Mark E. Fornace, Aneesh Acharya, Georgios Artavanis, Johannes Stegmaier, Alexandre Cunha, and Niles A. Pierce. Third-generation in situ hybridization chain reaction: multiplexed, quantitative, sensitive, versatile, robust. Development, 145(12), June 2018.

[78] Emily Kuehn, David S. Clausen, Ryan W. Null, Bria M. Metzger, Amy D. Willis, and B. Duygu Özpolat. Segment number threshold determines juvenile onset of germline cluster expansion in platynereis dumerilii. Journal of Experimental Zoology Part B: Molecular and Developmental Evolution, 338(4):225240, November 2021.

[79] Rodrigo Ibarra-García-Padilla, Aubrey Gaylon Adam Howard, Eileen Willey Singleton, and Rosa Anna Uribe. A protocol for whole-mount immuno-coupled hybridization chain reaction (wichcr) in zebrafish embryos and larvae. STAR Protocols, 2(3):100709, September 2021.

[80] Miguel A Moreno-Mateos, Charles E Vejnar, Jean-Denis Beaudoin, Juan P Fernandez, Emily K Mis, Mustafa K Khokha, and Antonio J Giraldez. Crisprscan: designing highly efficient sgrnas for crispr-cas9 targeting in vivo. Nature Methods, 12(10):982988, August 2015.

[81] James A. Gagnon, Eivind Valen, Summer B. Thyme, Peng Huang, Laila Ahkmetova, Andrea Pauli, Tessa G. Montague, Steven Zimmerman, Constance Richter, and Alexander F. Schier. Efficient mutagenesis by cas9 protein-mediated oligonucleotide insertion and large-scale assessment of single-guide rnas. PLoS ONE, 9(5):e98186, May 2014.

[82] Charles E. Vejnar, Miguel A. Moreno-Mateos, Daniel Cifuentes, Ariel A. Bazzini, and Antonio J. Giraldez. Optimized crisprcas9 system for genome editing in zebrafish. Cold Spring Harbor Protocols, 2016(10):pdb.prot086850, October 2016.

